# Cardiolipin dynamics promote membrane remodeling by mitochondrial OPA1

**DOI:** 10.1101/2024.05.21.595226

**Authors:** Sirikrishna Thatavarthy, Luciano A. Abriata, Fernando Teixeira Pinto Meireles, Kelly E. Zuccaro, Akhil Gargey Iragavarapu, Gabriela May Sullivan, Frank R. Moss, Adam Frost, Matteo Dal Peraro, Halil Aydin

**Author notes:** These authors contributed equally to this work (Sirikrishna Thatavarthy, Luciano A. Abriata).

## Abstract

Cardiolipin (CL) is a mitochondria-specific phospholipid that forms heterotypic interactions with membrane-shaping proteins and regulates the dynamic remodeling and function of mitochondria. However, the precise mechanisms through which CL influences mitochondrial morphology are not well understood. In this study, employing molecular dynamics (MD) simulations, we determined that CL molecules extensively engage with the paddle domain (PD) of mitochondrial fusion protein Optic Atrophy 1 (OPA1), which controls membrane-shaping mechanisms. Structure-function analysis confirmed the interactions between CL and two conserved motifs of OPA1 at the membrane-binding sites. We further developed a bromine-labeled CL probe to enhance cryoEM contrast and characterized the structure of OPA1 assemblies bound to the CL-brominated lipid bilayers. Our images provide direct evidence of CL enrichment within the OPA1-binding leaflet. Last, we observed a decrease in membrane remodeling activity for OPA1 in lipid compositions with increasing concentrations of monolyso-cardiolipin (MLCL). This suggests that the partial replacement of CL by MLCL, as observed in Barth syndrome-associated mutations of the tafazzin phospholipid transacylase, alters the malleability of the membrane and compromises proper remodeling. Together, these data provide insights into how biological membranes regulate the mechanisms governing mitochondrial homeostasis.

## Introduction

The proper spatial and temporal organization of organelles underlies many cellular processes, ranging from division and differentiation to apoptosis and communication^1^. Within a cell, mitochondria are mainly organized into highly dynamic and interconnected networks, whose diverse functions are dependent on their complex structure and organization^2^. Mitochondria are double-membrane bound organelles that consist of four major compartments: the outer membrane (OM), intermembrane space (IMS), inner membrane (IM), and matrix^3^. The mitochondrial IM folds inwards to form the organelle’s hallmark cristae membranes, which harbor the respiratory supercomplexes that produce ATP via oxidative phosphorylation (OXPHOS)^4^. In addition to their role in energy production, mitochondria are involved in the metabolism of amino acids, lipids, and nucleotides, transport of metabolites and ions, reactive oxygen species (ROS) production, and signaling^4,5^. The molecular regulation of mitochondrial architecture, which is controlled by the membrane-shaping lipids and proteins, is critical for tuning the activity of these key processes and preserving homeostasis^6–11^. Hence, mitochondrial function is intimately linked to dynamic changes in mitochondrial morphology and can influence human health and disease^12^.

The main lipid components of mitochondrial membranes are phospholipids^13,14^. Cardiolipin (CL) is a mitochondrion-specific phospholipid primarily located in the mitochondrial inner membrane (IM), where it accounts for ∼20% of the lipid content^13,14^. Characterized by a unique chemical structure consisting of a double glycerophosphate backbone and four fatty acyl chains^15^, CL undergoes maturation through biosynthesis and remodeling processes catalyzed by different enzymes within mitochondria^16–19^. Mature CL molecules interact with and regulate several pivotal proteins in mitochondria, including those involved in the regulation of mitochondrial morphology^20–26^. Aberrant CL content, structure, and localization result in mitochondrial defects and cellular dysfunction, leading to the development of cardiovascular diseases^27^, impaired neuronal function ^28^, and neurodegeneration^29,30^. Barth syndrome, an X-linked disease conventionally characterized by dilated cardiomyopathy, skeletal myopathy, cyclic neutropenia, arrhythmias, growth retardation, and cognitive dysfunction, occurs in 1 in 300,000 to 400,000 births^31–33^. The predominant locus for this disorder has been mapped to the distal region of chromosome Xq28, which encodes the human tafazzin (TAZ)^32,34^. TAZ functions as a phospholipid transacylase, facilitating the transfer of acyl groups from phospholipids to monolyso-cardiolipin (MLCL) to generate mature CL species^16,35^. Mutations associated with Barth syndrome compromise TAZ function, resulting in alterations in CL level and molecular composition, along with defects in mitochondrial architecture and function^36–39^. Despite extensive research on the pathophysiology of abnormal CL acyl composition arising from defective remodeling in cellular models, the molecular mechanisms connecting MLCL accumulation and protein function remain poorly understood.

CL plays an essential role in regulating the shape and stability of the mitochondrial IM by forming critical interactions with mitochondria-shaping proteins, determining the spatial identity and fitness of the organelle^21,40,41^. One such key protein involved in the modulation of mitochondrial architecture is optic atrophy 1 (OPA1), a mechano-chemical enzyme that catalyzes the fusion of mitochondrial IM, reorganizes dynamic cristae structure, and influences OXPHOS efficiency, apoptosis, reactive oxygen species production, and mtDNA maintenance^42–46^. In humans, the OPA1 precursor gives rise to eight isoforms, all of which are directed to the mitochondrial intermembrane space (IMS)^47^. Subsequently, divergent proteolytic mechanisms first cleave the mitochondrial-targeting sequence (MTS) to produce the long form (L-OPA1), which is N-terminally anchored to the inner membrane (IM), followed by the generation of the short form (S-OPA1) devoid of the transmembrane (TM) domain^48,49^. Both L-OPA1 and S-OPA1 assemble into oligomers and participate in membrane remodeling, and are essential for maintaining mitochondrial organization^49^. All OPA1 variants and proteoforms assemble into higher-order oligomers in the presence of CL-containing membranes^10,11,50–52^. This CL-OPA1 interaction is sufficient to activate membrane fusion and uphold cristae structural integrity, thus highlighting the regulatory role of CL in mitochondrial remodeling and function.

Understanding the precise molecular interactions between key mitochondrial proteins and CL within intact membranes has not been possible because single-phase fluid bilayers are generally thought to lack a structured pattern at the nanoscale. Hence, our understanding of molecular mechanisms connecting CL and mitochondrial protein function remains incomplete. To address this challenge and investigate CL’s functional role within the structural organization of mitochondrial membranes, we conducted molecular dynamics (MD) simulations, performed biochemical studies, and devised a novel lipid labeling approach for CL localization in electron cryo-microscopy (cryoEM) maps. Our findings reveal how CL regulates the activity of mitochondria-shaping human OPA1 to maintain mitochondrial homeostasis and provide a molecular explanation for the mechanisms underlying the disruptive effects of MLCL accumulation on mitochondrial membrane dynamics.

## Results

### Microsecond CG MD simulations reveal OPA1 membrane binding sites

In a recent study, we reported the cryoEM structures of human S-OPA1 helical assemblies bound to CL-containing lipid tubes^10^. These findings unveiled the architecture of assembled OPA1 and large structural arrangements potentially involved in catalyzing mitochondrial IM fusion. However, the mechanistic understanding of how OPA1 molecules selectively engage with CL-enriched membranes to modulate mitochondrial morphology remains unclear. To understand the molecular basis of CL-dependent mitochondrial remodeling, we conducted coarse-grained molecular dynamics (CG MD) simulations on microsecond time scales using S-OPA1 tetramers and lipid bilayers mimicking the composition of the mitochondrial IM^13,14^. While human S-OPA1 forms micron-scale helical filaments upon membrane binding^10^, simulating filamentous assemblies over relevant timescales proved not feasible due to their large size. Instead, we focused on tetrameric arrangements of S-OPA1 proteins, which encompass all key assembly interfaces and are tractable through multi-microsecond sampling at the coarse-grained level. To initiate simulations, we extracted four different tetrameric subassemblies (tetramers 1 to 4) of S-OPA1 models representing various oligomeric and functional states of the protein from the cryoEM helical reconstruction of the membrane-bound human S-OPA1 polymer (Figs. 1a-b and Supplementary Fig. S1). These assemblies were manually positioned with their membrane-interacting surface proximal to, but not fully inserted into, a model membrane composed of a mixture of MARTINI lipids POPC, POPE, and CL(18:1)_4_ at a ratio of ∼4:4:2, respectively (Supplementary Figs. S2a-c). The protein-membrane systems were parameterized with MARTINI22P, reaching close to 1 million beads that represent ∼10 million atoms in each system. We simulated each of these 4 tetrameric systems for over 8 µs to assess whether and how the tetramers engage with the membranes. Tetramer 1, representative of the conserved crisscross association of dynamin superfamily proteins^53^, was simulated in three independent replicas to maximize sampling and allow for the most accurate comparisons (Fig. 1c).

**Figure 1.**
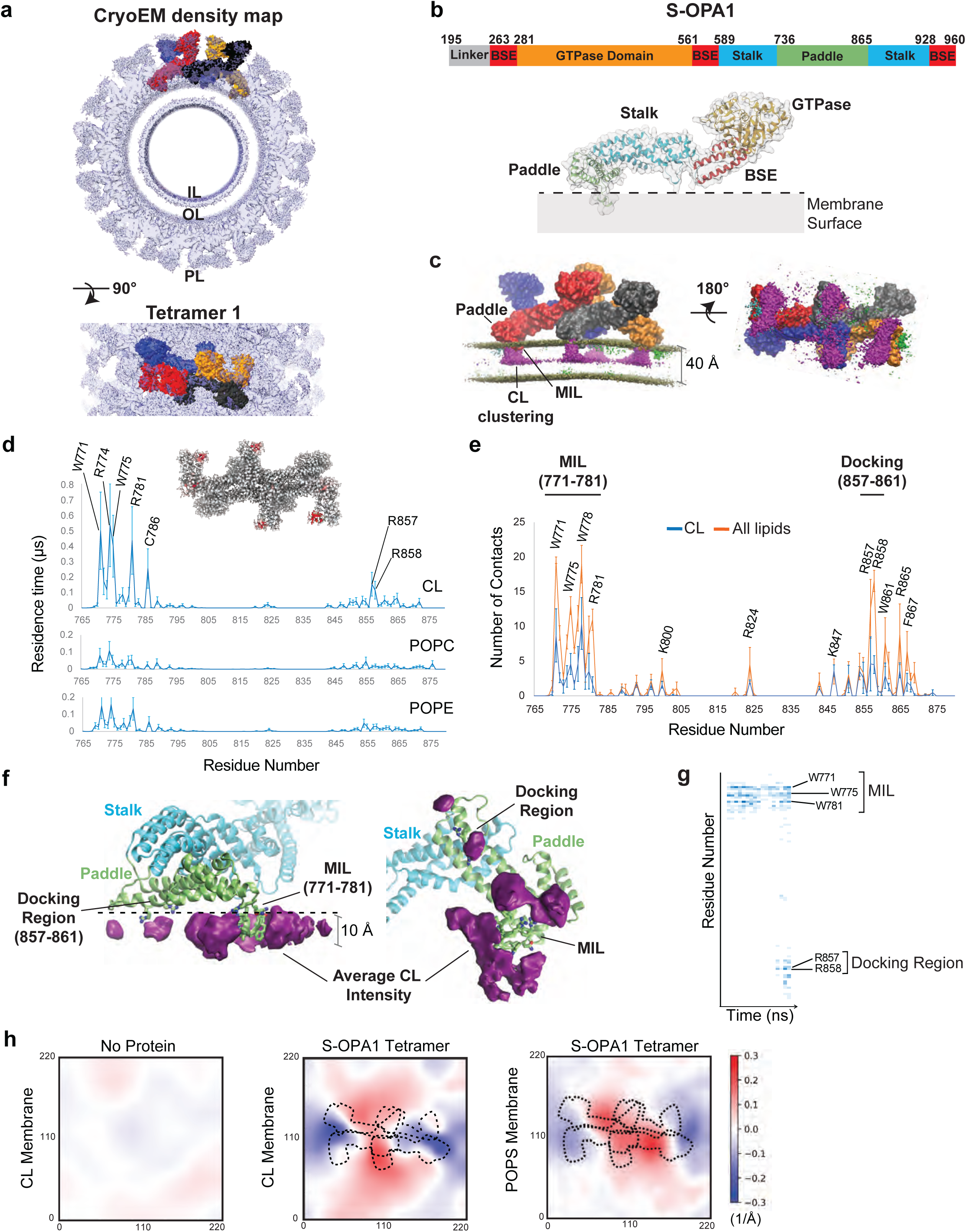
Interactions between lipids and S-OPA1 residues and changes in membrane topology. **(a)** The S-OPA1 tetramer model was extracted from the cryoEM structure of the membrane-bound S-OPA1 polymer in membrane-proximal conformation (PDB ID: 8CT1) and fitted into the cryoEM density map (EMDB ID: 26977). Each subunit of the tetramer is shown in surface representation and a different color, matching those in panel (c). IL, Inner Leaflet; OL, Outer Leaflet; PL, Protein Layer. **(b)** Structural organization of human S-OPA1. The primary sequence of S-OPA1 contains four domains: bundle signaling element (BSE), GTPase, stalk, and paddle. The extracted ribbon model for S-OPA1 monomer is colored in orange (GTPase), red (BSE), blue (stalk), and green (PD) to highlight the domain organization of S-OPA1, and the surface is depicted as semi-transparent solid density. This color scheme is used throughout the manuscript, except otherwise noted. **(c)** Tetramer 1 after docking onto a membrane containing 17% CL, 44% POPC, and 39% POPE via unbiased CG MD simulations. Each OPA1 subunit is shown in surface representation with the colors defined in **(a)**. Note the curvature experienced by the membrane in the direction of tubulation with OPA1 remaining outside, quantified in panel **h**. The average densities for all beads that make up the CG MD simulation lipids are shown in magenta (CL), green (POPC), and cyan (POPE). Despite being minoritarian in composition, the CL molecules cluster very strongly at the protein-membrane contact sites. Bottom view of the S-OPA1 tetramer simulated on CL-enriched membranes is shown on the right. **(d)** Residence times for contacts between protein and lipid beads are shown for CL, POPE, and POPC and are calculated for all four subunits in each of the three CG MD simulation replicates for tetramer 1. The inset above the plot for CL shows an OPA1 tetramer in surface representation with the residence times mapped as shades increasing from grey to red, to highlight the positions of those residues with longer residence times. **(e)** The average number of protein-lipid contacts per residue was calculated using the last 300 ns of the AA MD simulations in 3 independent replicas. **(f)** CL density averaged from AA MD simulations also shows that this lipid clusters strongly at OPA1-membrane contact sites, especially around the MIL (residues 771 to 781) and the docking region (residues 857 to 861). The simulations were set up by using a monomeric S-OPA1 model extracted from tetramer 1 and placing it onto membranes with the same lipid composition, as in the CG simulations; however, CL(18:2)_4_ was used for AA MD simulations as this is the main CL species in both cardiac and skeletal muscle tissues. **(g)** The heat map was generated using one of the replicas that shows strong binding to the membrane and displays the number of membrane contacts for S-OPA1 residues over time (nanoseconds). For more details about CG and AA MD simulations, please see Methods. **(h)** Membrane deformation analysis in CG MD simulations. Membrane bending activity of the S-OPA1 tetramer 1 measured in CG MD simulations with a membrane patch containing 44% POPC, 39% POPE, and 17% CL or POPS. The three graphs correspond to top views of the membranes, color-coded by the replica- and trajectory-averaged (last 4µs out of 8µs) deformations along the membrane normal, measured in 1/Å. The images (from left to right) show the CL membrane without protein, the CL membrane with S-OPA1 tetramer, and the POPS membrane with S-OPA1 tetramer. Red and blue colors indicate membrane pulling and pushing in the direction of the membrane normal, respectively. The comparison of the three different membrane deformation analyses indicate that the S-OPA1 tetramer was able to bend membranes containing both CL and POPS, but the extent of deformation is decreased in the presence of POPS (fewer areas of deep blue and deep red). The membrane pulling in POPS-containing membranes peaks at ∼0.3 Å^-1^, while membrane pushing only reaches to -0.2 Å^-1^ in one region and is even weaker in others. The dashed lines correspond to an overlay of the position of the OPA1 tetramer.

Upon visual inspection, all four subunits of tetramer 1 exhibit clear and strong membrane binding within the first few hundred nanoseconds. In this timescale, the highly conserved membrane-inserting loop (MIL) region (^771^WKKRWLYWKNR^781^) of the paddle domain (PD) inserts into the lipid bilayer, firmly anchoring the assembly tightly onto the membrane (Fig. 1c, and Supplementary Figs. S2a-c and S5). A second highly conserved site within the PD (^857^RRGFY^861^), which we refer to hereafter as the “docking region”, also interacts with membranes, albeit peripherally. The docking region does not embed in the membrane but remains stably bound throughout the simulation, as quantified below through lipid-protein contact residence times (Supplementary Figs. S2a-c and S5). To address potential biases that may arise due to the initial proximity of S-OPA1 tetramers to the membrane (∼6 Å), we conducted a supplementary set of CG MD simulations. In this series, a single subunit of S-OPA1 tetramer 1 was initially positioned 60 Å away from the model membrane (Supplementary Fig. S2b). Across all five replicas, the S-OPA1 monomer eventually encountered the membrane. In four of the replicas, extended interactions between S-OPA1 and the bilayer were observed.

#### CL molecules cluster at OPA1 membrane binding sites in simulations

Upon encountering the membrane surface, the positively charged residues within the MIL region formed the initial charge-charge contacts with the negatively charged headgroups of CL molecules, which was followed by rapid engagement of key tryptophan residues with the membrane and the insertion of the MIL into the bilayer (Fig. 1d). The docking region then formed peripheral interactions with the membrane lipids, positioning the positively charged membrane-facing surface of the PD onto the bilayer. At this stage, the local lipid composition of the membrane patch near the protein contact sites remained unchanged. Concurrently with the insertion of the MIL into the membrane, CL rapidly localized at the protein-membrane contact sites and the number of sidechain-CL interactions increased to ∼50% of total contacts, reaching a higher average density compared to POPC and POPE despite its lower concentration in the simulated membrane (Figs. 1c-d). While POPC and POPE transiently interact with S-OPA1 at the membrane contact sites, CL molecules establish much stronger interactions, demonstrated by lipid contact residence times 5 to 10 times longer than the other phospholipids (Fig. 1d). Importantly, control CG MD simulations on the membrane without protein did not exhibit aggregation or phase separation of CL molecules, suggesting that S-OPA1 interactions with the lipid bilayer are the trigger for the recruitment of CL molecules to the protein-membrane contact sites.

Analysis of the residence times for contacts between protein residues and CL molecules revealed extensive engagement of CL with two specific motifs: MIL residues W771, K772, K773, R774, W775, W778, and R781 and R857 and R858 residues located in the docking region (Fig. 1d). Mutating some of the MIL and docking region residues to alanine within the same CG MD system abolished the membrane binding activity of the tetrameric subassemblies in simulations, and the assemblies remained disengaged in solution while retaining their quaternary structure (Supplementary Fig. S2d). These findings closely mirror our previous observations in experimentally determined structural models. Finally, to confirm the specificity of OPA1-CL interactions, we substituted CL with another negatively charged phospholipid, POPS, in the simulated membrane. These experiments showed short contact residence times for POPS and weaker interactions with OPA1 residues compared to CL (Supplementary Fig. S3a-b). Together, our data suggest that OPA1-membrane interactions are controlled by the two CL binding motifs within the PD of the protein.

#### AA MD simulations confirm local CL accumulation near protein contact sites

While the CG MD simulations already suggest that the positively charged beads of the lysine and arginine residues of the MIL and docking regions engage in contact with the negatively charged beads of CL head group and tryptophan residues of the MIL interact with the hydrophobic tails of lipids, we examined these interactions in greater detail through all-atom (AA) MD simulations. We first extracted one of the subunits of S-OPA1 tetramer 1 and embedded it in a membrane using the membrane-monomer orientations and interactions retrieved from the CG MD simulations. We utilized two separate membrane patches containing two different CL molecular species, CL(18:1)_4_ and CL(18:2)_4_, and ran three replicas of each system for ∼1 µs to facilitate free, unbiased exploration of the protein-lipid contacts. Similar to the CG MD simulations, the AA MD simulations revealed local accumulation of CL around the protein-membrane contact sites in all three replicas of each system, and we did not observe a difference between CL(18:1)_4_- and CL(18:2)_4_-containing membranes. CL molecules formed constant interactions in particular with the key CL binding motifs of the MIL and docking regions (Figs. 1e-g and Supplementary Fig. S3c-d). While the CG MD simulations displayed high density for the beads of the headgroup and acyl chains of CL, spanning across the OPA1-binding leaflet, the AA MD simulations indicate that only the headgroups contributed to the density in the same leaflet. These differences likely arise from the high flexibility of the CL tails, a feature only captured in the AA MD simulations.

Despite being only 20% of the membrane composition in all the AA MD simulations, CL accounts for nearly half of the protein-membrane contacts with the membrane-facing residues of the PD (Figs. 1e-g and Supplementary Fig. S3c-d). The key electrostatic interactions between CL and OPA1 are mediated by K772 and R781 of the MIL, R857 and R858 of the docking region, and other critical membrane interface residues, including K800, R824, K847, and R865, within the PD. R857 and R858 residues continue to interact completely peripherally with the CL phosphates via their guanidinium groups on the bilayer surface (Figs. 1e-g and Supplementary Fig. S3c-d). This data shows the preference of positively charged residues located at the membrane interface of S-OPA1 for specific interactions with the negatively charged headgroups of CL molecules, thereby recruiting them to the protein-membrane contact sites. However, there is a greater tendency for the CL molecules to be present in the vicinity of the CL binding motifs (Figs. 1f-g). Analysis of the last 100 ns of each AA MD trajectory reveals an average of ∼2 CL molecules in contact with the R857 and R858 residues of the docking region and ∼4 CL molecules around the MIL residues, suggesting that CL molecules are particularly enriched around the MIL motif of the PD (Figs. 1f-g and Supplementary Figs. S3c-d).

The AA MD simulations also revealed enriched hydrophobic contacts between CL and S-OPA1 PD near the MIL region. Following the initial protein-membrane contacts, predominantly driven by charge-charge interactions at the solvent-membrane interface, the membrane insertion of the MIL is facilitated by the indole rings of MIL residues W771, W775, and W778. Upon insertion, the tryptophan sidechains become vertically embedded into the spaces in-between lipids, causing the helix comprising the MIL to lodge deep into the membrane by ∼10 Å, equivalent to ∼25% of the bilayer thickness (Fig. 1f). Mechanistically, the insertion exposes MIL residues to the hydrophobic core of the membrane to facilitate direct interactions with the lipid tails. Interestingly, the three tryptophan residues exhibit less selectivity for the hydrophobic acyl chains of CL upon membrane insertion, forming similar interactions with the acyl chains of POPC and POPE. These findings indicate that charge-charge interactions facilitate the clustering of CL molecules in the vicinity of the PD. Once in close proximity, the CL tails interact with hydrophobic side chains of the MIL, further stabilizing S-OPA1 subunits on the membrane. Collectively, our computational findings demonstrate that the main structural element driving OPA1 activation on CL-containing membranes is the MIL motif, as it facilitates direct binding to CL headgroup and acyl chains, while the docking region plays a lesser role in OPA1-membrane dynamics.

#### S-OPA1 tetramer is capable of bending membranes

In the presence of CL-containing lipid vesicles, S-OPA1 molecules become activated rapidly and polymerize into membrane-remodeling filaments. This process marks the initial step in reshaping the mitochondrial IM, yet the precise stoichiometry of the OPA1 machinery required for initiating local membrane bending remains unknown. As a member of the dynamin superfamily, OPA1 functions through the oligomerization of its monomeric, dimeric, or tetrameric basic building blocks into rings or helices to remodel membranes in cells^53^. Consistent with this notion, our CG MD simulations using S-OPA1 tetramers demonstrate membrane bending in a direction conducive to ring formation of OPA1 proteins on the outer side of the formed tubule (Fig. 1h and Supplementary Fig. S4). Quantitatively, all four tetrameric subassemblies induce positive curvature protruding towards the protein, averaging up to ±0.3 Å^-1^ throughout the simulations at specific points where the membrane contacts the protein (Fig. 1h and Supplementary Fig. S4). For comparison, we measured a control membrane without protein and determined an average fluctuation of ±0.03 Å^-1^ (Fig. 1h). The most curved snapshots of the simulation display local curvature radii ranging from 20 nm to 50 nm, a range consistent with the ∼19 nm inner lumen diameter observed in our cryoEM structure of the S-OPA1 polymer wrapped around a membrane tube. A lower curvature radius indicates stronger membrane bending, and the deformations observed in CG MD simulations are likely limited by the strong lateral membrane pressure acting through periodic cells, preventing bending of the lipid bilayer. Additionally, S-OPA1 tetramer was able to bend POPS-containing membranes, albeit to a lesser extent (Fig. 1h). Similar bending induced by the protein also takes place with the other tetramers, as presented later in the manuscript.

#### Saturation of CL acyl chains hinders S-OPA1 activity

To test our computational models of OPA1 in biochemical assays, we employed an *in vitro* reconstitution assay using purified S-OPA1 and liposomes prepared with various lipid compositions (Supplementary Figs. S5a-b and Supplementary Table 2). We reconstituted human S-OPA1 WT samples onto lipid bilayers and quantified the membrane binding and remodeling activity of the protein by using co-sedimentation assays and negative-stain transmission electron microscopy (TEM). Consistent with our previous findings, analysis of the reconstitution assays revealed that S-OPA1 molecules fail to bind and polymerize on lipid vesicles when CL is omitted from lipid compositions (Supplementary Fig. S6). Similarly, substitution of CL with negatively charged phospholipids POPG and POPS impaired S-OPA1’s ability to remodel membranes (Supplementary Fig. S6). Although we observed some remodeling with POPS-containing liposomes, the total remodeled area was decreased by more than 95% when compared to the experiments with CL-enriched membranes (Supplementary Fig. S6c-d). To investigate the role of CL acyl chain composition on S-OPA1-mediated membrane organization, we comprehensively characterized five different molecular species of CL *in vitro.* We prepared liposomes with two saturated (CL(16:0)_4_ and CL(18:0)_4_) and three unsaturated (CL(16:0)_2_-(18:1)_2_, CL(18:1)_4_, and CL(18:2)_4_) CL species and tested their effect on S-OPA1 activity. We observed the highest amount of membrane binding and remodeling with CL(18:2)_4_, which is the most abundant CL species in cardiac and skeletal muscle cells (Fig. 2 and Supplementary Fig. S7). While we detected comparable S-OPA1 membrane binding activity with the CL(16:0)_2_-(18:1)_2_ and CL(18:1)_4_, this activity decreased by ∼70% with liposomes containing CL(16:0)_4_ and CL(18:0)_4_ (Figs. 2a-b). Next, S-OPA1-mediated membrane remodeling was measured in reconstitution assays. After 4hrs of reaction, S-OPA1 was able to form higher-order assemblies in the presence of CL(16:0)_2_-(18:1)_2_, CL(18:1)_4_, and CL(18:2)_4_, whereas CL(16:0)_4_ and CL(18:0)_4_ hindered the ability of S-OPA1 to assemble into stable elongated filaments, indicating a clear preference for unsaturated CL species over saturated species for remodeling (Fig. 2c-e and Supplementary Fig. S7). Together, these findings suggest that CL acyl chain composition is critical for the proper assembly and function of OPA1 on lipid membranes. Since CL(18:1)_4_ and CL(18:2)_4_ are found to be the dominating CL species in various tissues, we conducted the subsequent experiments using these two molecular species of CL.

**Figure 2.**
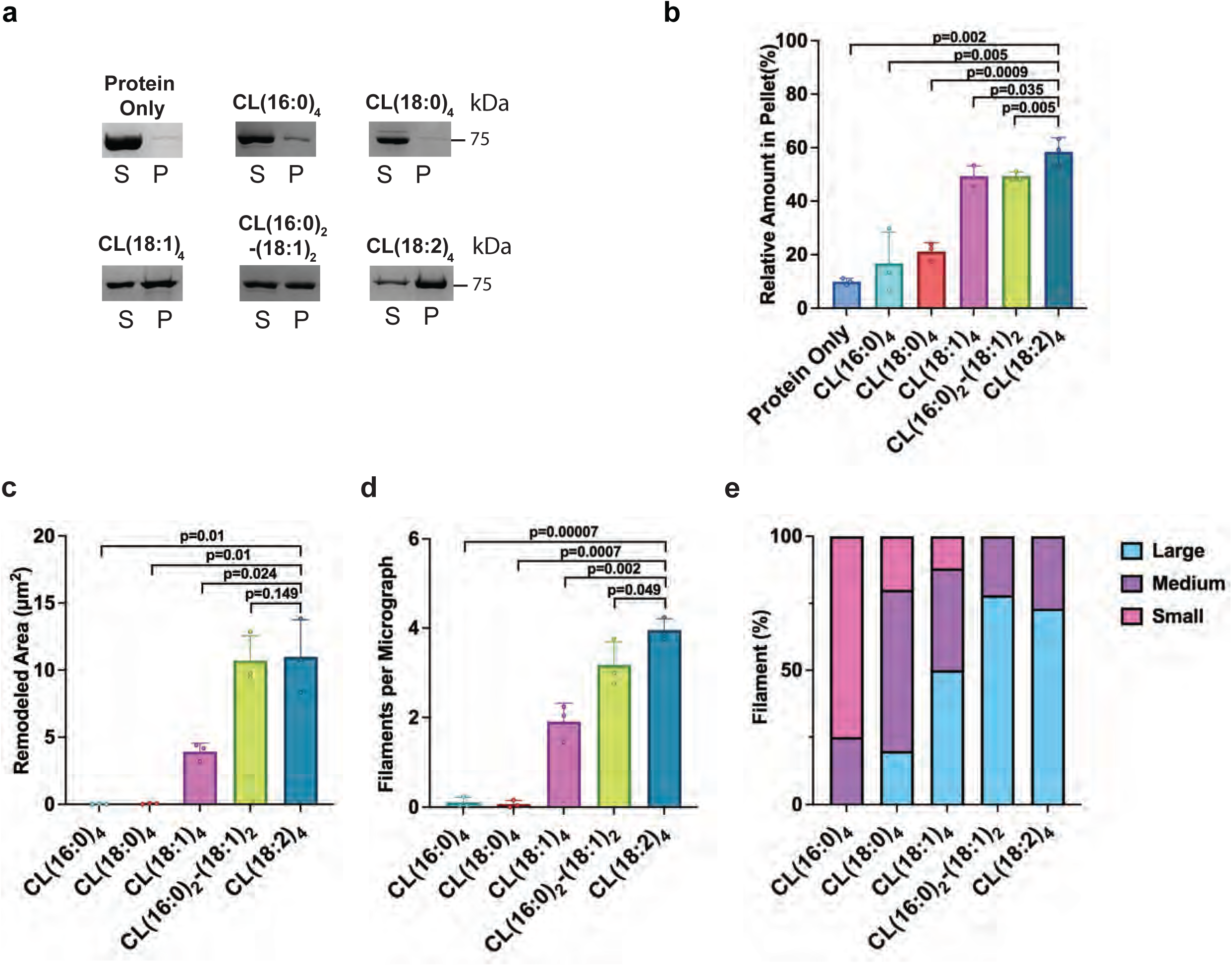
Membrane binding and remodeling experiments with different CL Species. **(a)** Co-sedimentation assays of S-OPA1 WT were performed using liposomes containing five different CL species: CL(16:0)_4_, CL(18:0)_4_, CL(18:1)_4_, CL(16:0)_2_-(18:1)_2_, and CL(18:2)_4_. All liposomes share a common composition of 45% POPC, 22% POPE, and 8% L-PI, with the remaining 25% composed of the respective CL species. Supernatant and pellet samples were collected after centrifugation, subjected to SDS-PAGE, and quantified using ImageJ. S, supernatant. P, pellet. **(b)** The assays were performed in triplicate, and the binding activity was quantified from the pellet fractions. Statistical significance was assessed using a two-tailed unpaired Welch’s t-test based on *n*=3 independent experiments comparing binding across various CL-containing liposome compositions. *P*-values are represented on the graph, and *p*<0.05 is considered to be statistically significant. **(c)** Remodeled area (μm^2^) was calculated by measuring the surface area of membrane tubules. Each lipid composition was tested in three independent reconstitution assays, and negative-stain TEM imaging was performed on each replicate. The total remodeled area was calculated per micrograph and a total of 75 negative stain micrographs were analyzed per lipid composition. **(d)** S-OPA1 filaments were counted in each micrograph and plotted against the corresponding lipid composition. **(c, d)** Error bars represent the standard error of the mean (s.e.m.). Statistical significance was assessed using a two-tailed unpaired Welch’s *t*-test with *n*=3 independent experiments. *P*-values are annotated on the graph, and *p*<0.05 is considered statistically significant. **(e)** Quantification of filament morphology per liposome composition. Each remodeled filament was counted and assigned to one of the three classes: large (cyan), medium (purple), or small (pink). Filament sizes were defined as small at 0-0.01 μm^2^, medium for 0.01-0.05 μm^2^, and large if >0.05 μm^2^. The percentage of each class is calculated per liposome composition and is shown as a stacked bar graph.

#### The two CL binding motifs contribute to the membrane remodeling activity of S-OPA1

To verify the functional relevance of the two CL binding motifs, we created, recombinantly expressed, and purified three mutant constructs, as well as the WT construct. An alanine point mutation was introduced to the positively charged R858 residue within the docking region motif. A second mutant construct was generated to replace the charged and hydrophobic MIL region motif (^771^WKKRWLYWKNR^781^) with a polyalanine stretch, and an R858A-MIL polyalanine double-mutant was employed to show the importance of both CL-binding motifs. While the individual R858A and MIL polyalanine mutants reduced the membrane binding activity by ∼12% and ∼54%, respectively, the double mutant led to a ∼60% decrease in binding. Together, both mutants exhibited reduced membrane binding activity on CL-enriched liposomes compared to the WT (Figs. 3a-b). We then determined that R858A and MIL polyalanine mutations impair S-OPA1’s ability to form ordered assemblies and remodel CL-containing lipid membranes *in vitro* (Figs. 3c-f). Notably, the R858A mutant diminished the total remodeled area by approximately 80% and generated more small size filaments, while the MIL region mutant and the double-mutant resulted in abrogation of remodeling function *in vitro* (Figs. 3c-f). Our biochemical data indicate that the two conserved motifs form stable interactions with CL molecules, ensuring the proper assembly of OPA1 polymers on the membrane and promoting mitochondrial morphology remodeling.

**Figure 3.**
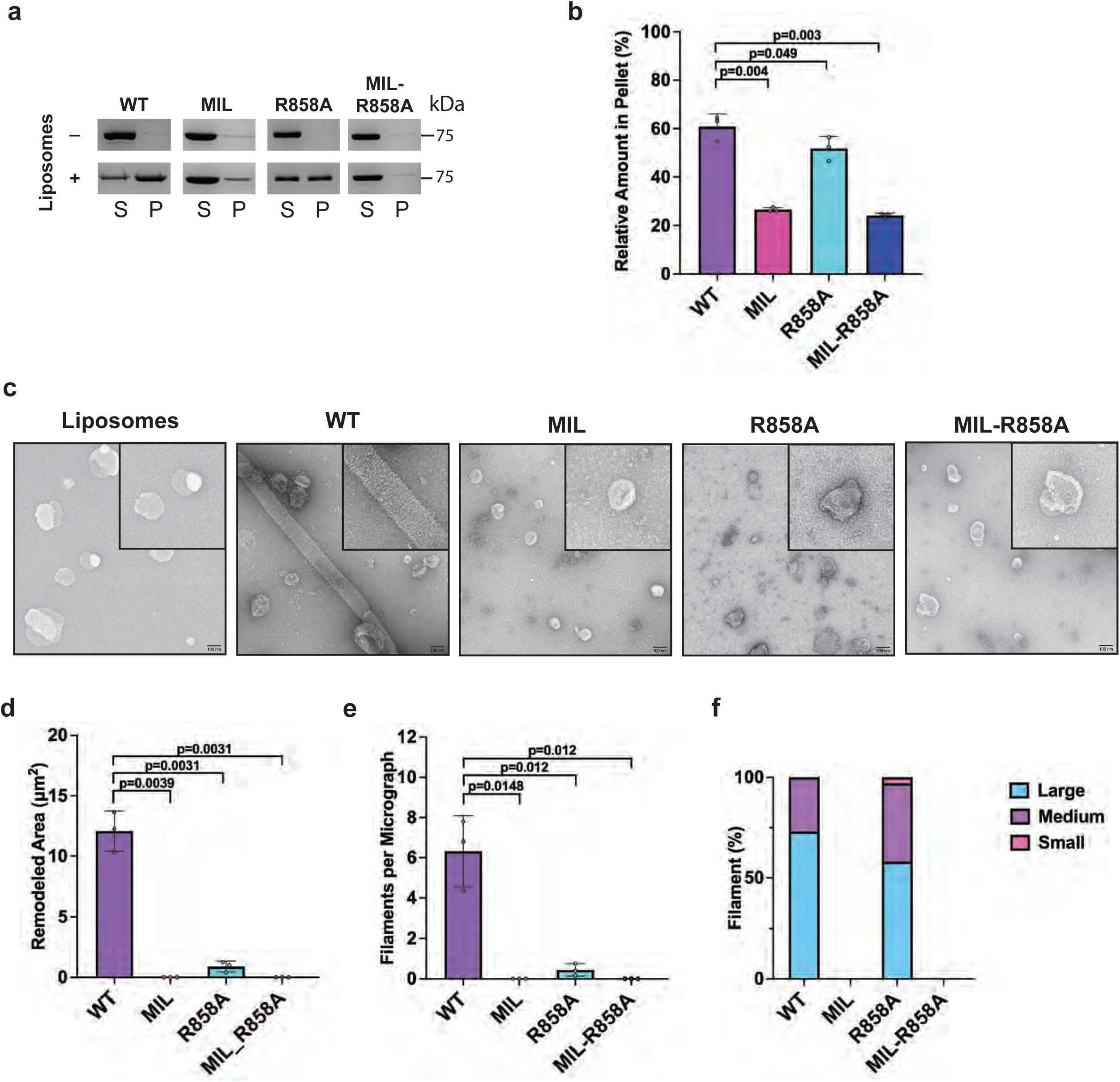
Characterization of key membrane interface residues. **(a)** The SDS-PAGE analysis of the co-sedimentation assays with S-OPA1 WT, MIL, R858A, and MIL-R858 double-mutant in the presence and absence of CL-containing liposomes. S, supernatant. P, pellet. A lipid mixture of 45% POPC, 22% POPE, 8% L-PI, and 25% CL(18:2)_4_ was used to prepare liposomes. **(b)** Supernatant and pellet samples were collected after centrifugation, subjected to SDS-PAGE, and quantified using ImageJ. The assays were performed in triplicate, and the binding activity was quantified from the pellet fractions. Statistical significance was assessed using a two-tailed unpaired Welch’s t-test based on *n*=3 independent experiments comparing S-OPA1 WT to mutant constructs. *P*-values are represented on the graph, and *p*<0.05 is considered to be statistically significant. **(c)** The reconstitution assays for WT and mutant S-OPA1 samples were visualized by negative-stain TEM. Scale bar is 100 nm. **(d)** Remodeled area (μm^2^) was calculated by measuring the surface area of membrane tubules. WT and mutant samples were tested in three independent reconstitution assays, and negative stain TEM imaging was performed on each replicate. The total tubulated area was calculated in each micrograph and a total of 75 micrographs were analyzed per sample. **(e)** S-OPA1 filaments per micrograph were counted and plotted against the corresponding sample. **(d, e)** Error bars represent the standard error of the mean (s.e.m.). Statistical significance was assessed using a two-tailed unpaired Welch’s *t*-test with *n*=3 independent experiments. *P*-values are annotated on the graph, and *p*<0.05 is considered statistically significant. **(f)** Quantification of filament morphology per liposome composition. Each remodeled filament was counted and assigned to one of the three classes: large, medium, or small. Filament sizes were defined as small at 0-0.01 μm^2^ (pink), medium for 0.01-0.05 μm^2^ (purple), and large if >0.05 μm^2^ (cyan). The percentage of each class is calculated per sample and is shown as a stacked bar graph.

#### Structure of S-OPA1 assembly bound to membranes containing contrast-enhancing probes

To experimentally determine CL localization in intact lipid membranes, we synthesized CL with bromine atoms added to the unsaturated fatty acyl chains. This modification capitalizes on the lipophilicity, steric, and enhanced electron scattering properties of Br, and results in dibrominated lipid tails mimicking unsaturated tails that facilitate the determination of the position of CL molecules within the structural organization of lipid bilayers (Fig. 4a and Supplementary Fig. S8)^54,55^. We reconstituted human S-OPA1 onto lipid bilayers (both vesicles and nanotubes) containing brominated CL and learned that S-OPA1 can self-organize into higher-order structures on these bilayers and induce the protrusions of narrow lipid tubes (Supplementary Fig. S9a). This observation indicates that the brominated CL, which yields stronger electron scattering, exhibits similar membrane packing properties and behaves indistinguishably from unsaturated phospholipids *in vitro*. To measure and model the interactions between CL and OPA1, we prepared samples for cryoEM and recorded images of S-OPA1 filament segments bound to brominated CL-containing lipid tubes using a 300-kV Krios cryoEM microscope (Supplementary Fig. S10 and Supplementary Table 1). Image segments were first aligned and averaged to obtain *ab initio* 3D reconstructions, followed by 3D classification to generate well-ordered subsets using the Relion software (Supplementary Figs. S10 and S11)^56^. However, the membrane-bound assemblies exhibited slightly variable tubule diameters, hindering coherent inter-tube averaging and resulting in multiple conformational classes. To address this variability, we performed 3D classification without alignment and identified filament segments with nearly uniform diameters. The best quality maps were then refined to obtain a sub-nanometer reconstruction of S-OPA1 polymer bound to brominated lipid membranes (Supplementary Fig. S11). The density map distinctly delineates two components corresponding to the protein coat and lipid bilayer, with numerous S-OPA1 subunits forming a spiraling homomeric filament on membranes (Fig. 4 and Supplementary Fig. S10). This reconstruction represents the membrane-proximal conformation of the OPA1 assembly, wherein the PD is docked on the membrane surface and the MIL is embedded in the lipid bilayer (Figs. 4b-d).

**Figure 4.**
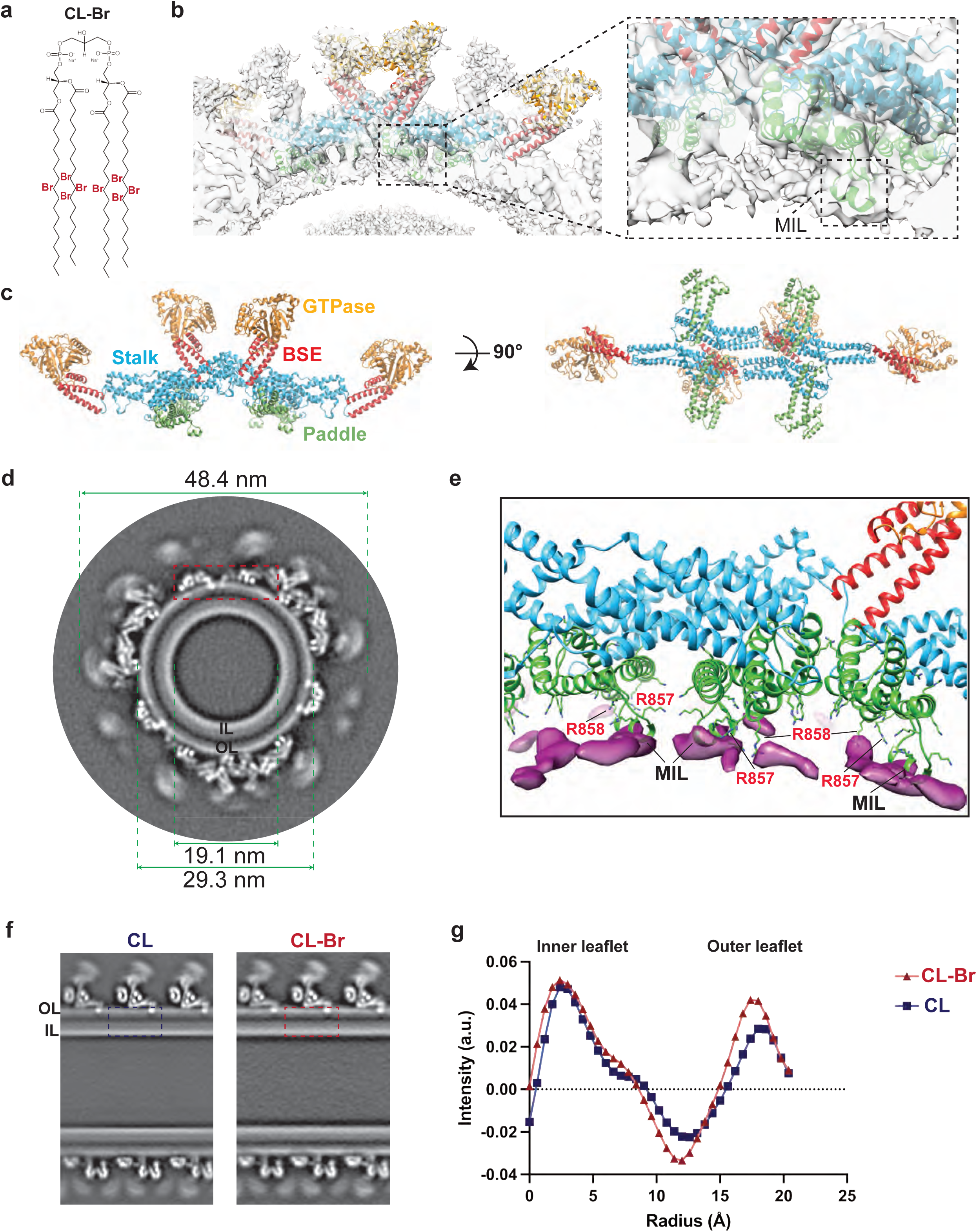
CryoEM reconstruction of human S-OPA1 bound to CL-Br-enriched membranes. **(a)** Structure of tetrabrominated CL. **(b)** The side view for surface representation and the corresponding ribbon diagram of the S-OPA1 tetramer 1 oriented into the cryoEM density map. The inset window shows the close-up view of the paddle domain (green) and conserved MIL region interacting with membranes. **(c)** Tetrameric ribbon model of the S-OPA1 bound to CL-Br membranes. The four structural domains are colored as follows: GTPase (orange), BSE (red), Stalk (blue), and Paddle (green). **(d)** A gray-scale slice of the cryoEM 3D reconstruction of the membrane-bound S-OPA1 filament. The red rectangle indicates the position of the magnified view shown in panel E. **(e)** The difference map calculated from brominated and native protein-lipid reconstructions shows additional densities (magenta) located near the PDs (green) of S-OPA1. **(f)** Gray-scale slices of native and brominated membranes along the helical axis. The cryoEM reconstruction of S-OPA1 assembly bound to CL-Br-containing membranes shows CL enrichment in the outer leaflet. **(g)** Radial profiles from the cryoEM 3D reconstructions indicate the location of the surplus signals attributed to halogen scattering. The red box in panel A indicates the region used in intensity analysis.

The final 3D reconstruction of the membrane-bound S-OPA1 polymer reveals an outer diameter of 48.4 nm and an inner lumen diameter of 19.1 nm (Fig. 4d). It exhibits a three-start helical structure with a rise of 7.69 Å and a twist of 128.642 degrees, with minimal intersubunit connectivity arising from the low packing density of the S-OPA1 lattice (Figs. 4b-c, Supplementary Figs. S10 and S11, and Supplementary Table 1). The cryoEM density map achieved sufficient resolution to unambiguously assign the orientation of the S-OPA1 domains. While the bundle-signaling element (BSE), stalk, and PD could be resolved, the distal GTPase domains that are not interacting with the lipid bilayer were at the lower local resolution, indicating the dynamic nature and conformational flexibility in the membrane-bound state (Supplementary Fig. S10d). Nonetheless, leveraging this reconstruction and prior structural knowledge enabled us to build precise molecular models of S-OPA1 tetramers bound to brominated lipid membranes with an overall resolution of 6.4 Å (Fig. 4 and Supplementary Figs. S10 and S11). A comparison of membrane-bound OPA1 models from native and brominated liposomes showed highly similar structures with a root-mean-square deviation (RMSD) of only 0.78 Å over 698 Cα atoms of the protein (Supplementary Fig. S10e). These findings collectively indicate that bromine labeling of CL acyl chains does not induce notable structural changes in how OPA1 assembles on lipid membranes.

#### CryoEM structure shows CL enrichment in the OPA1-bound outer leaflet

To detect the position of CL molecules in the reconstructions, we investigated the membrane layer of the experimental density map for focal enrichment of CL-Br. Initially, we normalized the pixel value distributions to the S-OPA1 intensity from radial averages and obtained horizontal and vertical slices of brominated and non-brominated reconstructions (Figs. 4d-f). The density map confirmed that the docking and the MIL regions of the PD are positioned to make direct interactions with CL molecules in the lipid bilayer (Figs. 4d-e). Comparing the Coulombic potentials from the resulting 3D maps of unlabeled versus labeled membrane tubes, we located the surplus signals attributable to halogen scattering in the outer leaflet (Figs. 4f-g). Although CL-Br enrichment in the OPA1-bound leaflet was observed, the bilayers remained compositionally heterogeneous with CL-Br distributed throughout the bilayers as observed in the difference map between the CL membrane and CL-Br membrane (Figs. 4f-g and Supplementary Fig. S10f). In CG MD simulations, CL molecules form frequent microsecond-timescale interactions with the membrane-facing residues of OPA1 PD. Thus, the combined experimental and computational data imply a dynamic nature of CL within lipid bilayers, which is critical for OPA1 anchoring. These results collectively demonstrate that halogenated lipids scatter electrons strongly, enabling the quantitative localization of surplus scattering in our cryoEM maps for estimating the changes in CL concentration within each leaflet. Additionally, identifying overall enrichment of CL in the outer leaflet with cryoEM supports our CG MD data, providing us a platform to further explore the mechanics governing mitochondrial membrane reshaping at higher resolution.

#### MLCL forms similar clusters near protein-membrane contact sites

Next, we investigated whether the accumulation of MLCL in lipid bilayers affects OPA1’s ability to reshape membranes and control the dynamic architecture of mitochondria. We utilized CG MD simulations with the same four S-OPA1 tetramers and assessed how the replacement of CL by MLCL affects OPA1’s interactions with membranes. All parameters and the overall setup remained identical to the previous CG MD simulations, except for the substitution of CL with MLCL. After >8µs of simulations, all replicas of the trajectories for the four models revealed membrane binding and clustering of MLCL around the same CL binding motifs. Additionally, they exhibited similar protein-lipid interaction profiles and residence times compared to CL contacts.

To quantitatively analyze the MLCL-protein interactions, we measured the residence times of MLCL in the presence of S-OPA1 tetramers and compared them to the CL residence times. Within the uncertainty of the sampling in the CG MD simulations, residence times for protein-lipid contacts were similar between MLCL and CL, and both residence times were larger than those observed for negatively charged POPS in control experiments (Supplementary Fig. S3a). Likewise, MLCL- and CL-containing lipid bilayers displayed a similar number of protein-lipid contacts per residue in AA MD simulations, despite MLCL containing one fewer acyl chain (Supplementary Fig. S3b). Collectively, our MD simulations at CG and AA resolutions revealed no major differences in how human OPA1 interacts with membranes containing CL or MLCL.

#### MLCL accumulation impairs OPA1’s ability to bend and remodel membranes

Despite the similar protein-lipid interactions and residence times observed with MLCL and CL in the bilayers, the deformation experienced by the MLCL-containing membranes is substantially weaker in the simulations, especially for assemblies mediated by the conserved crisscross association of S-OPA1 monomers (tetramers 1 and 3) (Fig. 5a and Supplementary Fig. S4). The simulations suggest that differences in the spontaneous curvature and other material properties of CL and MLCL are critical for OPA1-mediated membrane remodeling, with CL enabling membrane shape plasticity and bilayer deformation upon protein binding. Furthermore, S-OPA1 can also warp POPS-containing membranes to some extent (Fig. 1h and Supplementary Fig. S6), indicating that the intrinsic mechanical properties of lipid bilayers may play an important role in governing OPA1’s ability to remodel membranes.

**Figure 5.**
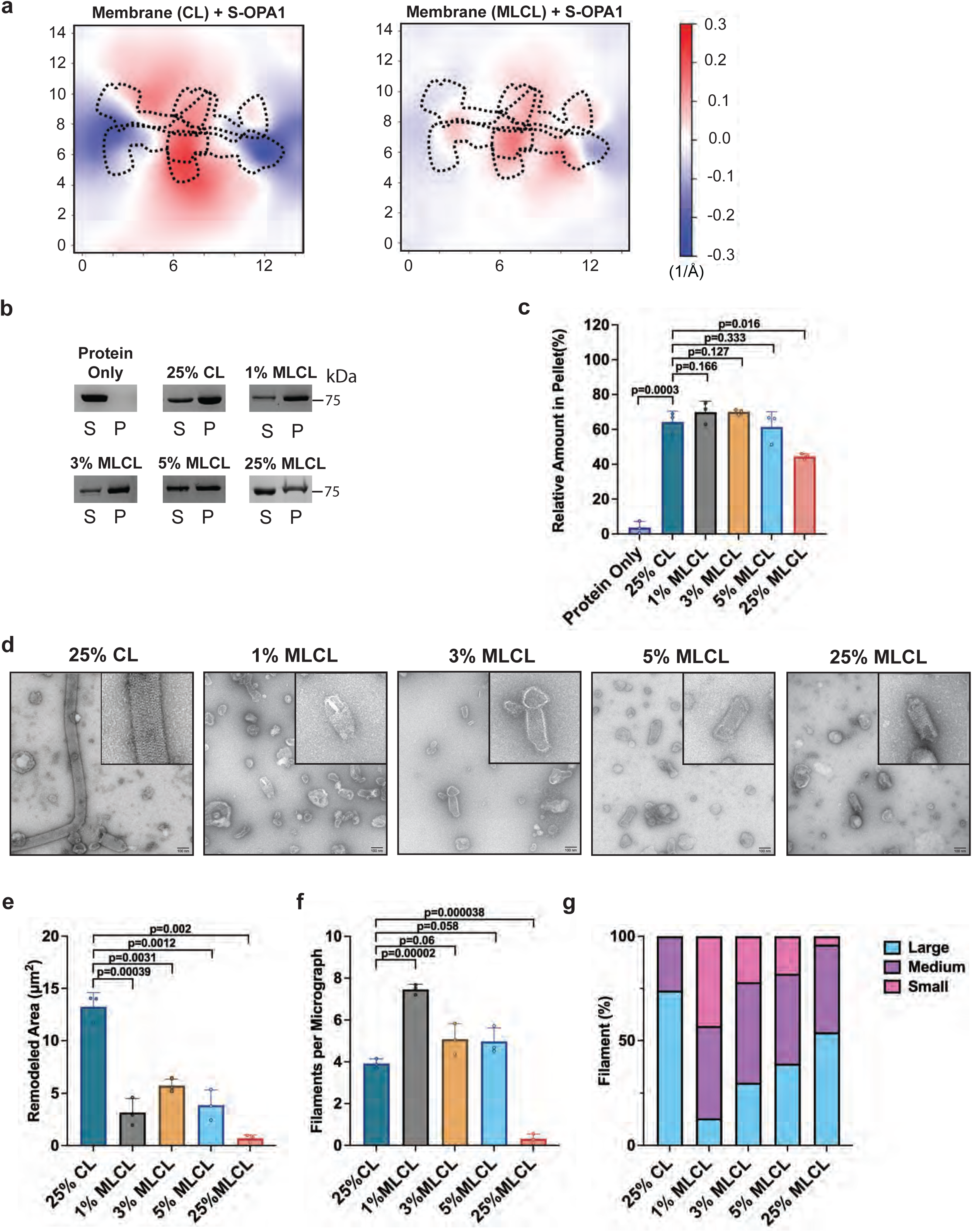
S-OPA1 interactions with MLCL-containing membranes. **(a)** Membrane deformation calculations are shown for one of the three independent replicas using S-OPA1 tetramer 1 and model membranes containing either CL (left) or MLCL (right). Red and blue colors indicate membrane pulling and pushing in the direction of z, respectively. The membrane deformation activity of tetramer 1, particularly its ability to push down on the sides, is reduced in the presence of MLCL. **(b)** The co-sedimentation assays with S-OPA1 WT and liposomes containing CL and increasing molar ratios of MLCL. A lipid mixture of 45% POPC, 22% POPE, 8% L-PI, and 25% CL and/or MLCL was used to prepare five different liposomes. The CL:MLCL ratios used for liposome preparations as follows: 25% CL; 24% CL and 1% MLCL; 22% CL and 3% MLCL; 20% CL and 5% MLCL; and 25% MLCL. Supernatant and pellet samples from the co-sedimentation assays were harvested after centrifugation and analyzed by SDS-PAGE. **(c)** The assays were performed in triplicate and gel images were quantified by ImageJ. The binding activity of S-OPA1 WT was quantified from pellet fractions, and statistical significance was assessed using a two-tailed unpaired Welch’s t-test based on *n*=3 independent experiments, comparing CL-containing liposomes to those with increasing molar concentrations of MLCL. *P*-values are represented on the graph, *and p*<0.05 is considered to be statistically significant. **(d)** Representative negative-stain TEM images of reconstitution assays in the presence of CL- and MLCL-containing liposomes. Increasing molar ratios of MLCL alter the membrane remodeling activity of S-OPA1. Scale bars are 100 nm. **(e)** Remodeled area (μm^2^) was calculated by measuring the surface area of membrane tubules. Each lipid composition was tested in three independent reconstitution assays, and negative stain TEM imaging was performed on each replicate. The total remodeled area was calculated per micrograph and a total of 75 negative stain micrographs were analyzed per sample. **(f)** S-OPA1 filaments were counted in each micrograph and plotted against the corresponding lipid composition. **(e, f)** Error bars represent the standard error of the mean (s.e.m.). Statistical significance was assessed using a two-tailed unpaired Welch’s *t*-test with *n*=3 independent experiments. *P*-values are annotated on the graph, and *p*<0.05 is considered statistically significant. **(g)** Quantification of filament morphology per liposome composition. Each remodeled filament was counted and assigned to one of the three classes: large (cyan), medium (purple), or small (pink). Filament sizes were defined as small at 0-0.01 μm^2^, medium for 0.01-0.05 μm^2^, and large if >0.05 μm^2^. The percentage of each class is calculated per liposome composition and is shown as a stacked bar graph.

To experimentally probe the mechanistic basis of MLCL interactions with human OPA1, we performed co-sedimentation experiments with S-OPA1 and liposomes containing increasing concentrations of MLCL in place of CL (Supplementary Fig. S9b and Supplementary Table 2). Consistent with MD simulations, increasing concentrations of MLCL resulted in only very minor differences in OPA1’s ability to bind liposomes. We found that the presence of 1% to 5% of MLCL in liposomes resulted in similar membrane binding activity for S-OPA1 compared to liposomes containing 25% CL (Figs. 5b-c). Increasing MLCL concentration to 25% diminished S-OPA1’s ability to bind liposomes, resulting in a ∼30% decrease in membrane binding (Figs. 5b-c). The major difference in the chemical structure of MLCL is the absence of a single acyl chain compared to CL. The results of our MD simulations and co-sedimentation assays indicate that while the changes in the acyl chain content of CL molecular species do not affect the recruitment of OPA1 to the lipid membranes at low molar concentrations, the full replacement of CL by MLCL decreases the membrane binding activity of the protein.

Following this, we investigated the impact of MLCL on the membrane remodeling activity of S-OPA1. We reconstituted S-OPA1 with CL- and MLCL-containing liposomes and monitored the protein’s oligomerization and membrane remodeling activity using negative-stain TEM imaging (Fig. 5d). While the reconstitution of S-OPA1 on CL-containing liposomes resulted in the formation of higher-order protein assemblies and further tubulation of membranes, the replacement of CL by MLCL hindered the liposome remodeling activity of the protein and resulted in more than 90% decrease in total remodeled area (Figs. 5d-g). Lowering MLCL concentrations to 1 to 5% in the lipid compositions still reduced the membrane remodeling ability of S-OPA1 by more than 50% and the activity of the protein was not recovered. This suggests that even the lower molar concentrations of MLCL in lipid bilayers are enough to interrupt OPA1 polymerization on membranes, which is detrimental to OPA1-mediated mitochondrial remodeling (Figs. 5d-g). Finally, we reconstituted S-OPA1 on membrane nanotubes containing increasing concentrations of MLCL (1 to 10%) and determined that MLCL accumulation does not alter the formation of higher-order S-OPA1 assemblies on pre-curved membranes *in vitro* (Supplementary Figs. S9c-d). This suggests that the presence of MLCL increases the energy barrier required for local membrane bending.

Overall, while we did not observe notable reductions in how OPA1 binds CL- and MLCL-containing lipid bilayers, membrane remodeling experiments demonstrated that the presence of MLCL in lipid bilayers alters the remodeling activity of OPA1 machinery. Our findings suggest that MLCL accumulation changes the malleability of the lipid bilayer and restricts OPA1’s ability to remodel membranes. Hence, we postulate that OPA1 relies on the lipid composition of mitochondrial membranes for its activity, which is precisely tuned by the unique properties of CL and its enrichment at the OPA1-membrane contact sites.

In our proposed mechanism, OPA1 proteins are recruited to the membrane via specific interactions with CL molecules, which are randomly distributed throughout the membrane with potential pre-formed patches enriched in CL within the lipid bilayer. Upon protein binding, CL molecules rapidly localize to the outer leaflet of the bilayer near the two CL binding motifs, facilitating the formation of stable interactions between OPA1 and lipid bilayers. At these CL-rich contact sites, unique structural properties of CL and its stable interactions with key OPA1 residues allow for membrane deformation. Concomitantly, by leveraging these specific CL contacts, OPA1 proteins form higher-order assemblies required for the remodeling and the fusion of the mitochondrial IM. The partial replacement of CL by MLCL, even at low concentrations, makes the membrane less prone to bending, which in turn disrupts OPA1 activity, thereby hindering membrane remodeling mechanisms (Fig. 6).

**Figure 6.**
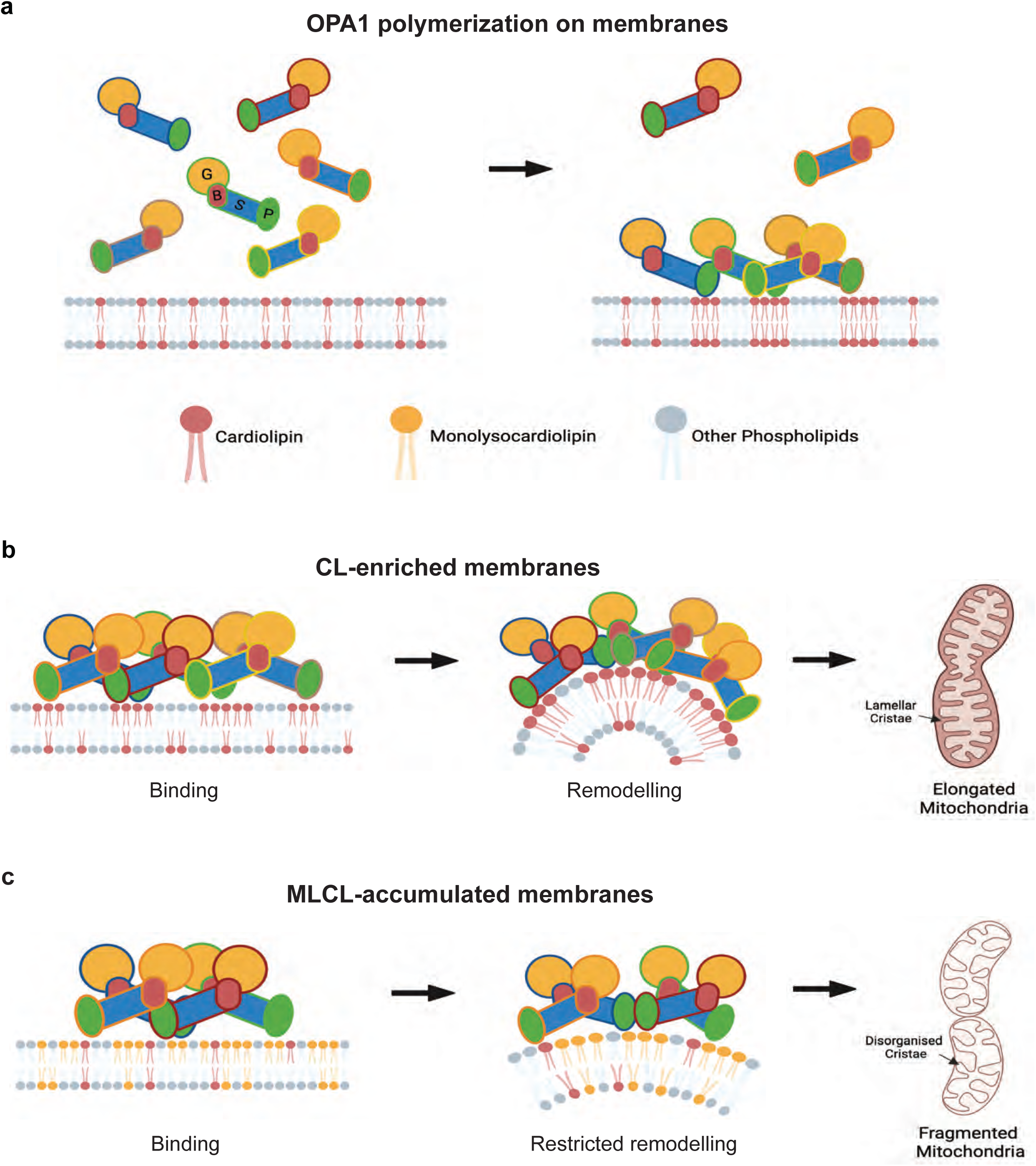
Proposed model of how CL controls mitochondrial remodeling. OPA1 interactions with randomly distributed CL molecules in membranes trigger the clustering of CL near protein-membrane contact sites and facilitate the remodeling of membranes. The accumulation of MLCL in membranes restricts the membrane remodeling activity of OPA1 and causes abnormalities in mitochondrial morphology.

## Discussion

Here, we performed multiple CG and AA MD simulations to explore the lipid-lipid and lipid-protein interactions driving OPA1-mediated mitochondrial remodeling. These simulations were set up with various OPA1 structures and the membrane bilayer mimicking the lipid composition of the mitochondrial IM. Dynamic models generated through these simulations revealed the CL dynamics within the bilayer, exhibiting enrichment in the outer leaflet and extensive contacts with OPA1 residues. Furthermore, our computational approach enabled us to accurately measure the lateral chemical organization and morphological changes in CL-enriched membranes that often occur at shorter time and length scales and are challenging to probe experimentally at the molecular level. These analyses led to the identification of two highly conserved binding motifs located at the MIL and docking regions of the PD, showcasing strong interactions with CL molecules within intact lipid membranes (Supplementary Figs. S3c-d and S5c). The CL binding motifs establish critical electrostatic and hydrophobic interactions with both the headgroup and acyl chains of CL, reminiscent of protein complexes observed in oxidative phosphorylation^23,24^, and thereby facilitate membrane remodeling. As anticipated, mutations of the key residues to alanine hindered the membrane binding and remodeling activity of OPA1 in our computational and biochemical assays. Note that our reconstitution assays and MD simulations offer an approximation of biological membranes. Moreover, our results align with previous studies, demonstrating that mutations to membrane-interacting residues of OPA1 result in fragmented mitochondrial morphology in living cells^10,11^. Overall, these findings allowed us to pinpoint specific lipid-protein interactions and understand how CL regulates the activity of OPA1 to maintain mitochondrial homeostasis.

CL is a pivotal regulatory lipid playing critical roles in various mitochondrial processes, including energy production, apoptosis, mitophagy, oxidative stress, and mitochondrial fusion and fission, which govern mitochondrial shape and function. Yet, for many of these cellular processes, it is currently unknown how the role of CL extends from maintaining membrane structure to an intimate association with mitochondrial proteins. Central to these processes is the intricate interplay between CL and the OPA1 protein that results in the initiation of OPA1-mediated mitochondrial membrane remodeling and maintenance of a healthy organellar network distributed throughout the cell. Thus, CL assumes a direct and regulatory role in the structural and functional remodeling of mitochondria.

The most distinguishing features of CL are its dimeric structure and polymorphic phase behavior^57–59^. CL and unsaturated phosphatidylethanolamine have the ability to organize into non-lamellar phases, particularly the hexagonal II phase, to promote local structures, such as negative curvatures, within membranes^16,33–36,39^. Previous studies reported that in the absence of divalent cations, CL prefers the lamellar phase at physiological temperature to form an extended bilayer. Ca^2+^ promotes tighter packing of CL headgroups and can induce a geometry that favors the hexagonal phase^57–59,65,66^. We found that CL is also highly dynamic and can diffuse near protein contact sites within microseconds throughout the simulated bilayer. Surprisingly, our simulations revealed that CL forms multiple short-lived interactions with the OPA1 side chains at the CL binding motifs rather than long-lived, stable interactions to modulate the formation of the OPA1 scaffold for membrane remodeling. Together, these findings suggest that both the polymorphic phase behavior and the rapid lateral diffusion of CL within the bilayer play a crucial role during membrane shape transitions.

Determining the functional role of CL within the structural framework of intact lipid bilayers poses challenges due to the heterogeneity and dynamic nature of mitochondrial membranes. Prior research has demonstrated that brominated and iodinated lipids behave similarly to their native counterparts and can serve as contrast-enhancing probes to delineate specific lipids within membranes^54,55^. Moreover, the utilization of bromo-substituents on aliphatic double bonds has a well-established history as fluorescence quenchers in model membranes^67–69^. By labeling CL with halogen atoms, which scatter electrons more strongly than acyl chains alone, we quantitatively located the surplus scattering in our cryoEM maps and estimated the concentration of CL within each leaflet. Our observations of the OPA1 structure bound to brominated membranes provide experimental evidence that CL is enriched in the OPA1-binding leaflet to promote the remodeling of mitochondrial membranes. We anticipate that this versatile tool will prove instrumental in determining how CL either activates or inhibits other key membrane-associated processes in the regulation of mitochondrial morphology and function.

On the other hand, human OPA1 forms several homo- and heterotypic interactions with proteins and lipids to modulate the dynamic remodeling of mitochondrial network^49,70–74^. In addition to its key interactions with CL, the interplay between L-OPA1 and S-OPA1 is crucial for controlling mitochondrial IM fusion and cristae dynamics. Previously, it was shown that L-OPA1 can tether and hemifuse bilayers but is unable to transition through the final step of pore opening or mediate low levels of pore opening^51^. Hence, optimal fusion requires a combination of L-OPA1 and S-OPA1 together, which synergistically catalyzes efficient and fast membrane pore opening^51,75^. Other studies reported that L-OPA1 alone does not have a notable fusion activity under normal cellular conditions but becomes more fusogenic under specific cell stress conditions^76^. Finally, L-OPA1 was shown to be sufficient to mediate fusion in cells lacking the proteases (YME1L and OMA1) responsible for generating S-OPA1^48,49^. It appears that the heterotypic OPA1 fusion machinery has a complex regulatory mechanism, and the relative contribution of L-versus S-OPA1 to membrane remodeling and fusion remains incompletely understood. L-OPA1 and S-OPA1 vary only by a short transmembrane helix (residues 88-109) and a linker region (residues 110-195). We postulate that the linker region provides sufficient structural freedom for membrane-anchored L-OPA1 to form higher-order structures analogous to those we observe for S-OPA1, and for L-OPA1 and S-OPA1 to co-assemble.

Barth syndrome (BTHS) stands as a significant X-linked cardiomyopathic disease characterized by perturbations of cardiolipin (CL) metabolism in mitochondria^31,32^. Despite its prevalence, the precise repercussions of altered lipid content underlying BTHS symptoms remain unclear. Here, we sought to determine how loss of CL content and accumulation of MLCL in mitochondrial membranes influence the activity of the key membrane remodeling enzyme, OPA1. Human OPA1 governs mitochondrial shape, cristae integrity, and functional output for a vast array of essential metabolic pathways and processes that determine cell function and fate. Initially, we investigated the impact of MLCL accumulation on OPA1-membrane interactions via MD simulations and measured the dynamics of the lipids throughout the bilayer to determine how MLCL molecules engage with OPA1 in the bilayer. Our simulations with CL-enriched membranes demonstrated that even though the CL molecules continuously diffuse throughout the membrane, they frequently associate with the two CL binding motifs. For instance, the interactions between the key binding motif residues W775 and R858 and CL persisted for 1 μs to 1.5 μs in residence times. We observed similar S-OPA1-membrane contacts when CL was replaced with MLCL in lipid compositions and measured comparable MLCL residence times for most of the MIL and docking region binding motif residues. This outcome is unsurprising, given that the main components mediating the initial protein-membrane interactions are the membrane-facing lysine and arginine residues of the PD and negatively charged glycerophosphate headgroups of CL and MLCL. The initial steps are followed by the MIL insertion in the membrane, which is mediated by the tryptophan residues within the MIL and their subsequent association with the hydrophobic acyl chains of phospholipids sharing similar physicochemical properties. However, the presence of MLCL hindered OPA1’s ability to bend membranes in MD simulations. Further probing of mechanistic links between MLCL and OPA1 activity through biochemical assays confirmed that the replacement of CL with MLCL reduces OPA1’s ability to remodel membranes. In these assays, S-OPA1 was able to bind and remodel the membranes that contained increasing MLCL concentrations, but the remodeling activity of the protein was decreased compared to the CL-enriched lipid bilayers. Interestingly, even partial replacement of CL with low molar concentrations of MLCL in membranes restricted the membrane-shaping activity of OPA1. Together, these findings indicate that MLCL accumulation could influence mitochondrial membrane properties, and the regulation of the mitochondrial IM morphology and associated cellular processes by human OPA1 depends on the lipid composition of the lipid bilayers.

Understanding the intrinsic mechanical properties of CL and MLCL, as well as their interactions with mitochondrial proteins, is critical for determining the molecular basis of pathologies associated with MLCL accumulation ^77^. Notably, CL is well known to both facilitate membrane bending to produce positive curvature and to partition into negatively curved regions^78–81^. The distinctive feature of MLCL is the absence of an acyl chain, which induces drastic differences in bilayer mechanical properties and curvature-dependent partitioning behavior^77–79^. More specifically, MLCL prefers a lamellar phase due to its cylindrical cone shape and induces less membrane curvature. The differences in the chemical structure could also influence the conformation of acyl tails and the accessibility of the headgroup due to hydrogen bonding with the additional hydroxyl group in MLCL, thereby affecting lipid conformations in the vicinity of MLCL. Hence, membranes containing increased concentrations of MLCL would be less malleable by membrane-shaping proteins, including human OPA1. Additionally, membrane binding and remodeling are interconnected processes occurring at different regulatory steps that modulate OPA1 activity. It is plausible that moderate differences in membrane binding could have drastic effects on the more energy-demanding membrane remodeling steps. Overall, these findings indicate that MLCL-containing membranes exhibit greater resistance to protein-mediated shape changes and provide insights into the disruptive effects of MLCL accumulation on mitochondrial membrane dynamics and function.

Despite the longstanding recognition of the significance of CL in modulating mitochondrial morphology and function, the precise mechanisms through which CL regulates these essential cellular machines, as well as the impact of the MLCL accumulation on mitochondrial membrane remodeling, remain unclear. Our findings highlight how CL molecules cluster near the membrane-binding surfaces of OPA1’s PD, engaging in charged and hydrophobic interactions with the conserved CL binding motifs to modulate the activity of the membrane-remodeling enzyme. Moreover, we describe how MLCL build-up in lipid membranes disrupts OPA1’s ability to remodel membranes, potentially playing an important role in the pathogenesis of inherited disorders, such as Barth Syndrome. These insights provide a critical structure-function foundation for understanding the mechanisms connecting CL and regulation of mitochondrial morphology, thus establishing a molecular basis for shaping the mitochondrial IM in health and disease.

## Materials and Methods

### Cloning, expression, and purification

The gene fragment corresponding to S-OPA1 (Addgene plasmid ID: 26047; residues 252-960) was subcloned into the pCA528 vector, incorporating an N-terminal 10X His tag followed by a SUMO solubility tag. S-OPA1 mutations were engineered using a modified QuickChange Mutagenesis protocol and confirmed through Sanger Sequencing. All S-OPA1 variant constructs were transformed into BL21 DE3-RIPL competent cells. A single colony from each transformation was inoculated into lysogeny broth (LB) media (100 ml) and was grown overnight at 37 °C with kanamycin (50 μg mL^-1^) and chloramphenicol (25 μg mL^-1^). The overnight culture (10 ml) was used to inoculate a 750 ml culture of ZYP-5052 auto-induction media and was grown at 37 °C until the optical density at 600 nm (OD_600_) reached a value between 0.6 and 0.8. At this point, the temperature was reduced to 18 °C within the shaker, and cultures continued to grow overnight for an additional 16 hours. Following the 16-hour induction period, the cells were harvested via centrifugation and stored at -80 °C.

The frozen bacterial pellets were thawed, resuspended with lysis buffer (50 mM HEPES-NaOH, pH 7.5, 500 mM NaCl, 20 mM imidazole, 5 mM MgCl_2_, 5 mM CHAPS (Anatrace), 5 mM 2-mercaptoethanol, 10% (v/v) glycerol) supplemented with 0.5% Triton X-100, 0.5 mg DNAseI, 1X EDTA-free complete protease inhibitor cocktail (Roche), and lysozyme. The cells were then lysed using an Emulsiflex C3 homogenizer or a probe sonicator (Qsonica Q500). To remove the cell debris, the lysate was centrifuged at 35,000 x g for 45 minutes at 4 °C. Meanwhile, a Ni-NTA (Qiagen) affinity column was equilibrated with lysis buffer. The supernatant was then filtered through a 0.45 μm membrane (Millipore) and transferred to a column, where it was incubated with the Ni-NTA beads on a roller for 1 hour at 4 °C. The column was then washed with 10 column volumes (CV) of lysis buffer followed by 10 CVs of high salt buffer (50 mM HEPES-NaOH, pH 7.5, 1 M NaCl, 20 mM imidazole, 5 mM MgCl_2_, 5 mM CHAPS, 5 mM 2-mercaptoethanol, and 10% (v/v) glycerol) and high imidazole buffer (50 mM HEPES-NaOH, pH 7.5, 500 mM NaCl, 80 mM imidazole, 5 mM MgCl_2_, 5 mM CHAPS, 5 mM 2-mercaptoethanol, and 10% (v/v) glycerol) washes. The sample was then eluted with 10 CVs of elution buffer (50 mM HEPES-NaOH, pH 7.5, 500 mM NaCl, 500 mM imidazole, 5 mM MgCl_2_, 5 mM CHAPS, 5 mM 2-mercaptoethanol, and 10% (v/v) glycerol). Following elution, the N-terminal 10XHis-SUMO tag was cleaved using the Ulp1 enzyme while dialyzing against the FPLC buffer (50 mM HEPES-NaOH, pH 7.5, 500 mM NaCl, 5 mM MgCl_2_, 5 mM CHAPS (Anatrace), 5 mM 2-mercaptoethanol, 10% (v/v) glycerol) at 4 °C. The digested protein samples were then concentrated with an Amicon Ultra (Millipore) concentrator (50 kDa MWCO) and subjected to further purification using a Superdex-200 16/60 column (Cytiva) equilibrated with the FPLC buffer for further purification. Pure fractions were pooled, concentrated to 2 mg mL^-1^, aliquoted, flash-frozen with liquid nitrogen, and stored at -80 °C for further use.

#### Preparation of lipid vesicles and nanotubes

All lipids, except for brominated cardiolipin (CL-Br), were purchased from Avanti Polar Lipids. Stock solutions were prepared by dissolving lipids in a chloroform, methanol, and water mixture (20:9:1, (v/v/v)) and stored in glass vials at -20 °C. The lipids in this study, 1-palmitoyl-2-oleoyl-glycero-3-phosphocholine (POPC), 1-palmitoyl-2-oleoyl-sn-glycero-3-phosphoethanolamine (POPE), 1-palmitoyl-2-oleoyl-sn-glycero-3-phospho-L-serine (POPS), 1-palmitoyl-2-oleoyl-sn-glycero-3-phospho-(1’-rac-glycerol) (POPG), L-α-lysophosphatidylinositol (Soy Lyso PI), 1’,3’-bis[1,2-dipalmitoyl-sn-glycero-3-phospho]-glycerol (CL (16:0)_4_), 1’,3’-bis[1,2-distearoyl-sn-glycero-3-phospho]-glycerol (CL (18:0)_4_), 1’,3’-bis[1-palmitoyl-2-oleoyl-sn-glycero-3-phospho]-glycerol (CL(16:0)_2_-(18:1)_2_), (1’,3’-bis[1,2-dioleoyl-sn-glycero-3-phospho]-glycerol (CL (18:1)_4_), heart CL (CL (18:2)_4_), and monolyso-cardiolipin (MLCL (94.6% (18:2)_3_ and 2.6% (18:1)_3_) were used as purchased without any modification. CL-Br was prepared by adapting previously described procedures for brominating alkenes in the lipids (*54*, *57*). Briefly, CL was dissolved in 5 mL of chloroform (ACS grade) and placed in a scintillation vial on ice. Liquid bromine, stoichiometric to the number of double bonds (Sigma Aldrich), was slowly added dropwise while the lipid solution was stirred on ice. The vial was then sealed and stirred on ice for 30 minutes in the dark. Solvent and excess bromine were removed under vacuum overnight, and the product was stored under a nitrogen atmosphere at -80 °C. The presence of bromine atoms on CL was confirmed with mass spectrometry (Supplementary Figs. S8a-b) and nuclear magnetic resonance (NMR) (Supplementary Figs. S8c-d). Before use, the CL-Br was warmed to room temperature and dissolved in chloroform to 5 mg mL^-1^. A lipid mixture of 45% POPC, 22% POPE, 8% PI, and 25% CL was used to prepare lipid vesicles mimicking the lipid composition of the mitochondrial inner membrane (IM)^82^. Conversely, CL-enriched lipid nanotubes were prepared using a lipid ratio of 90% D-galactosyl-(β)-1,1’N-nervonoyl-D-erythro-sphingosine (C24:1 Galactosyl(β) Ceramide, GalCer), and 10% CL or CL-Br. The other lipid compositions used in this study are provided in Supplementary Table 2. Vesicles and nanotubes were prepared following an established protocol^25^. Lipid stock solutions were warmed to room temperature for 15 minutes before they were mixed in a glass vial. The lipid mixtures were dried under a stream of nitrogen with rotation, and residual chloroform was further evaporated under vacuum overnight. The lipid film was resuspended in liposome buffer containing 20 mM HEPES-NaOH, pH 7.5, and 150 mM NaCl and rehydrated via vortexing. Unilamellar vesicles were prepared by extruding the rehydrated lipid film through a 50 nm pore-size polycarbonate membrane (Avanti), flash-frozen in liquid nitrogen, aliquoted, and stored at -80 °C. Lipid nanotubes were resuspended in liposome buffer with vortexing, sonicated with a bath-sonicator at 50 °C for 3-5 minutes until the lipid clumps were dissolved, and were used immediately.

#### Reconstitution assays and negative-stain transmission electron microscopy (TEM)

To set up reconstitution assays, the protein samples were further purified using a Superose 6 Increase 10/300 GL column (Cytiva) equilibrated with the reaction buffer (20 mM HEPES-NaOH, pH 7.5, 130 mM NaCl, 10 mM KCl, 2 mM MgCl_2_, 2 mM DTT, and 2% (v/v) glycerol). Purified samples (1.6 to 6 µM) were reconstituted with various liposomes and lipid nanotubes (Table S2) in the presence of 500 µM β,γ-methyleneguanosine 5′-triphosphate sodium salt (GMPPCP) for 4 hours at room temperature. Reconstituted samples were applied onto a glow-discharged metal mesh grid coated with carbon and stained with uranyl formate (0.75% w/v). Samples were visualized using negative-stain TEM. Images were collected on a Tecnai T12 Spirit TEM operating at 100 kV and equipped with an AMT 2k x 2k side-mounted CCD camera. Most images were recorded at a nominal magnification of 98,000x with a calibrated pixel size of 6.47 Å/pixel. Additional TEM micrographs were collected on a 120 kV Talos L120C TEM (Thermo Fisher Scientific) equipped with a Gatan 4k x 4k OneView camera at 17,500x magnification (pixel size, 8.4 Å/pixel). All experiments were performed in triplicate, and error bars indicate the standard error of the mean (s.e.m.).

#### Liposome co-sedimentation assays

Purified wild-type or mutant S-OPA1 proteins were buffer exchanged into the liposome buffer (20 mM HEPES-NaOH, pH 7.5, and 150 mM NaCl) using micro-spin desalting columns (Thermo Scientific Zeba). Equal volumes (25 μl) of protein (0.2 to 0.25 mg ml^-1^) and unilamellar vesicles (1.0 mg ml^-1^) were then mixed and incubated at room temperature for 30 min. After incubation, the samples were centrifuged at 55,000 rpm for 30 min at 20 °C using a TLA-120.2 rotor (Beckman Coulter). Reaction mixtures containing liposomes without CL were centrifuged at 85,000 rpm for 30 min to pellet proteoliposomes. The resulting supernatant and pellet fractions were subjected to SDS-PAGE analysis, and gel bands were quantified using ImageJ software^83^. Lipid compositions used in co-sedimentation assays are listed in Supplementary Table 2. All experiments were performed in triplicate, and error bars indicate the standard error of the mean (s.e.m.).

#### CryoEM grid preparation and data collection

For cryoEM, 6 μl of membrane reconstitution reaction containing lipid nanotubes with CL-Br was pipetted onto glow-discharged R1.2/1.3 200 copper mesh grids (Quantifold) in 100% humidity at 10 °C, incubated for 30 seconds, and blotted with a blot force of 0 for 4 seconds using a Vitrobot Mark IV (FEI). The grids were then plunge frozen into liquid ethane and stored under liquid nitrogen until imaged. Micrographs were acquired on a Titan Krios TEM (Thermo Fisher Scientific) operated at 300 kV and equipped with a K3 direct electron detector (Gatan) and a GIF Quantum energy filter (Gatan) with a slit width of 20 eV. SerialEM software^84^ was used for data acquisition. A total of 4,640 movie stacks were acquired with a defocus range of 0.5 to 1.5 μm at a nominal magnification of 105,000x, corresponding to a 0.417 Å/pixel in super-resolution mode. The movies were dose-fractionated into 118 frames with ∼0.55 e^−^ per Å^−2^ per frame and a total exposure time of 6 s, resulting in an accumulated dose of ∼65 e^−^Å^−2^ for each stack.

#### CryoEM data processing and 3D Reconstruction

Data processing procedure for membrane-bound OPA1 filaments was previously described^10^. Briefly, motion-corrected movies were imported into RELION 4.0^56^, and contrast transfer function (CTF) parameters were determined using CTFFIND4^85^. Following manual filament picking, a total of 233,341 segments were extracted from 4,640 micrographs with the data 2x binned by Fourier cropping (1.668 Å/pixel) and were subjected to multiple rounds of 2D and 3D classification, resulting in a subset of 11,469 particles. The resulting class was then subjected to 3D auto-refinement using a protein-only soft mask. Successive rounds of refinement were performed with higher resolution reference maps obtained after CTF and aberration refinements, which improved the map resolution to 6.7 Å. The helical parameters were refined to a rise of 7.69 Å and a twist of 128.642° per subunit. To further improve the signal-to-noise ratio, each independent half-map was segmented, resampled on a common grid, and summed according to the C2 symmetry axis of the OPA1 dimer using UCSF Chimera^86^. These summed unfiltered half maps were used during the post-processing step, yielding a final reconstruction at 6.4 Å resolution. The resolution of the final reconstructions was estimated by the Fourier Shell Correlation (FSC) between the two independent half maps at FSC=0.143. Resolution-dependent negative B-factors were applied to all final reconstructions for sharpening. Local resolution estimations were calculated using ResMap^87^. All cryo-EM data processing and analysis software was compiled and supported by the SBGrid Consortium^88^. An overview of cryo-EM data collection and image processing statistics was provided in Supplementary Fig. S11.

#### Model building, refinement, and validation

The cryoEM structure of membrane-bound S-OPA1 polymer (PDB ID: 8CT1) served as an initial reference for model building and refinement. Two distinct tetrameric models were extracted from the polymeric assembly and manually fitted into the density map using Chimera^86^. The tetrameric models underwent iterative refinement against the cryoEM map with global minimization, local grid search, and B factor refinement along with secondary structure, Ramachandran, and rotamer restraints to improve the model-map correlation coefficient using the phenix.real_space_refine tool in the PHENIX software package^89^. Further corrections to the models were made in Coot^90^ with torsion, planar peptide, and Ramachandran restraints. The quality of the model stereochemistry was validated by PHENIX and MolProbity^91^, and the model refinement and validation statistics are summarized in Supplementary Table 1. All structural figures were prepared in VMD^92^.

#### Molecular dynamics simulations

CryoEM structures of membrane-bound S-OPA1 polymers (PDB IDs: 8CT1 and 8CT9) were used as starting monomeric and tetrameric models for both CG and AA MD simulations. All MD simulation systems were set up using modules of the CHARMM-GUI server^93^, and then minimized, equilibrated, and run for production using standard CHARMM-GUI procedures, using Gromacs 2022^94^. Coarse-grained systems were prepared with MARTINI22p parameters and elastic networks^95,96^ for POPC, POPE, POPS, and CL, all inside the corresponding CHARMM-GUI module^97^, except for manual building and parametrization of MLCL membranes, which along with AA MD simulation files were kindly provided by Dr. Eric May, and described previously^79^. The dimensions of membranes for CG simulations were sized around 30nm x 30nm x 27nm, reaching around 1 million beads, which represent ∼10 million atoms in each system. MARTINI simulations were run for production at 303 K and 1 atm following minimization and thermal equilibration by using the standard procedures and parameters as provided by CHARMM-GUI. Briefly, a standard semi-isotropic Berendsen barostat was used for equilibration while progressively releasing positional restraints on the protein, and a standard Parrinello-Rahman semi-isotropic barostat was used without restraints. The integration timestep during production was 20 fs. The simulations were run for at least ∼8 µs, with the first few microseconds of trajectories removed for several analyses to obtain data computed on equilibrated systems. All CG simulations were run in 3 independent replicas.

Systems for atomistic simulations were prepared with CHARMM-GUI applying CHARMM36m parameters with the corresponding modified TIP3P^98^. The model membranes mimicking the lipid composition of the mitochondrial inner membrane (∼17% CL or MLCL, 44% POPC, and 39% POPE) were utilized in AA MD simulations and described previously^26^. System size was ∼20 nm x 20 nm x 20 nm and included over 600,000 particles. After minimization and equilibration to 303 K and 1 atm, production simulations were run using standard parameters as provided by CHARMM-GUI (standard semi-isotropic Berendsen barostat for equilibration while progressively releasing positional restraints on the protein and standard Parrinello-Rahman semi-isotropic barostat with a Nose-Hoover thermostat without restraints). PME electrostatics and a force-switch cut-off of 1 nm was applied, and an integration timestep of 2 fs was used during production for around 1 µs. Analyses were performed on the second half of the production phase to probe the equilibrated systems, a crucial step due to the slow convergence of lipid diffusion in AA MD simulations. All AA simulations were run in 3 independent replicas.

For all our CG and AA MD simulations, we used a total charge of -2 in the polar head of both CL and MLCL and utilized the corresponding parameters from MARTINI2.2p and CHARMM force fields as required. In CG MD simulations, where some 10-12 atoms are grouped and the exact degree of unsaturation is not resolved, only the CL(18:1)_4_ was used in simulations. The AA MD simulations were performed in the presence of both CL(18:1)_4_ and CL(18:2)_4_, and the production phase of each lipid composition and replicate was assessed separately. The pKa values for protein Lys and Arg sidechains were analyzed by PROPKA and predicted to be all in their charged forms, and were modeled accordingly. MD trajectories were visually inspected in VMD^92^ and analyzed with freely available tools and packages, detailed as follows. Atom-atom and bead-bead contacts were computed with tools from the PDB manipulation suite (https://lucianoabriata.altervista.org/pdbms/), applying 4 Å cutoffs on non-hydrogen atoms in atomistic simulations and 8 Å cutoffs for beads in CG simulations. Minimal distances between proteins and membranes were also measured with tools from the PDB manipulation suite. Residue-wise residence times per lipid were computed with the PyLipID package^99^ using standard settings. Volumes describing atom or bead densities were computed and visualized in VMD using the VolMap plugin, considering all non-hydrogen atoms when analyzing AA simulations and all beads when analyzing CG simulations. Membrane deformation was computed with the MembraneCurvature plugin for MDAnalysis, with standard settings and x,y grids of 14x14 tiles for tetramers 1 and 3 and 18x18 tiles for tetramers 2 and 4, which required a larger membrane surface. The trajectories subject to membrane deformation analysis were processed with Gromacs’ trjconv command, which centers the protein in the membrane by wrapping the frames and removes any translation and rotation on the plane.

### Statistical Analysis

Liposome co-sedimentation assays were performed in triplicate; the supernatant and pellet of each replicate were run on an SDS-PAGE gel, and the relative amount of protein was quantified via ImageJ. These values were converted to percentages, and only the pellet values were used for statistical analysis. An unpaired two-tailed Welch’s t-test was performed to compare the binding of protein across various experiments. Reconstitution assays were also performed in triplicate, and each replicate was quantified independently. Statistical analysis using unpaired two-tailed Welch’s test was carried out to compare cardiolipin with other lipids, as well as S-OPA1 WT and mutants. For both assays, p values <0.05 were considered statistically significant.

## Acknowledgments

The authors gratefully acknowledge NYU Langone Microscopy Laboratory (RRID: SCR_017934), especially Dr. Feng-Xia (Alice) Liang and Joseph Sall, for their support and assistance in this work. This shared resource is partially supported by the Cancer Center Support Grant P30CA016087 to the Laura and Isaac Perlmutter Cancer Center at NYU Langone Health. We would also like to thank Garry Morgan and Sarah Zimmermann at the EM Services Core Facility of the University of Colorado Boulder for electron microscopy training and support; the Shared Instrument Pool (SIP) core facility (RRID: SCR_018986) of the Department of Biochemistry at the University of Colorado Boulder for the use of the shared research instrumentation infrastructure; Dr. Annette Erbse for assistance with biophysical instruments and support; Lucero Castorena Casillas for carefully reading and editing the manuscript; Dr. Eric May for sharing files for MD simulation analysis.

## Author Contributions

S.T., K.E.Z., A.G.I., and G.M.S. performed cloning, mutagenesis, biochemical and biophysical characterizations, and carried out electron microscopy imaging and image analysis. F.R.M. synthesized brominated cardiolipin for cryoEM experiments and assisted with data collection. S.T., K.E.Z., G.M.S., and F.R.M. prepared liposomes for both biochemical and biophysical experiments. H.A. determined the cryoEM structures, and H.A. and K.E.Z. conducted model building, refinement, and validation of the cryoEM structures. L.A.A., F.T.P.M., and M.D.P. performed and analyzed the molecular dynamics simulations. All authors analyzed the data, discussed the results, and wrote the manuscript.

---

**Supplementary Figure 1.**
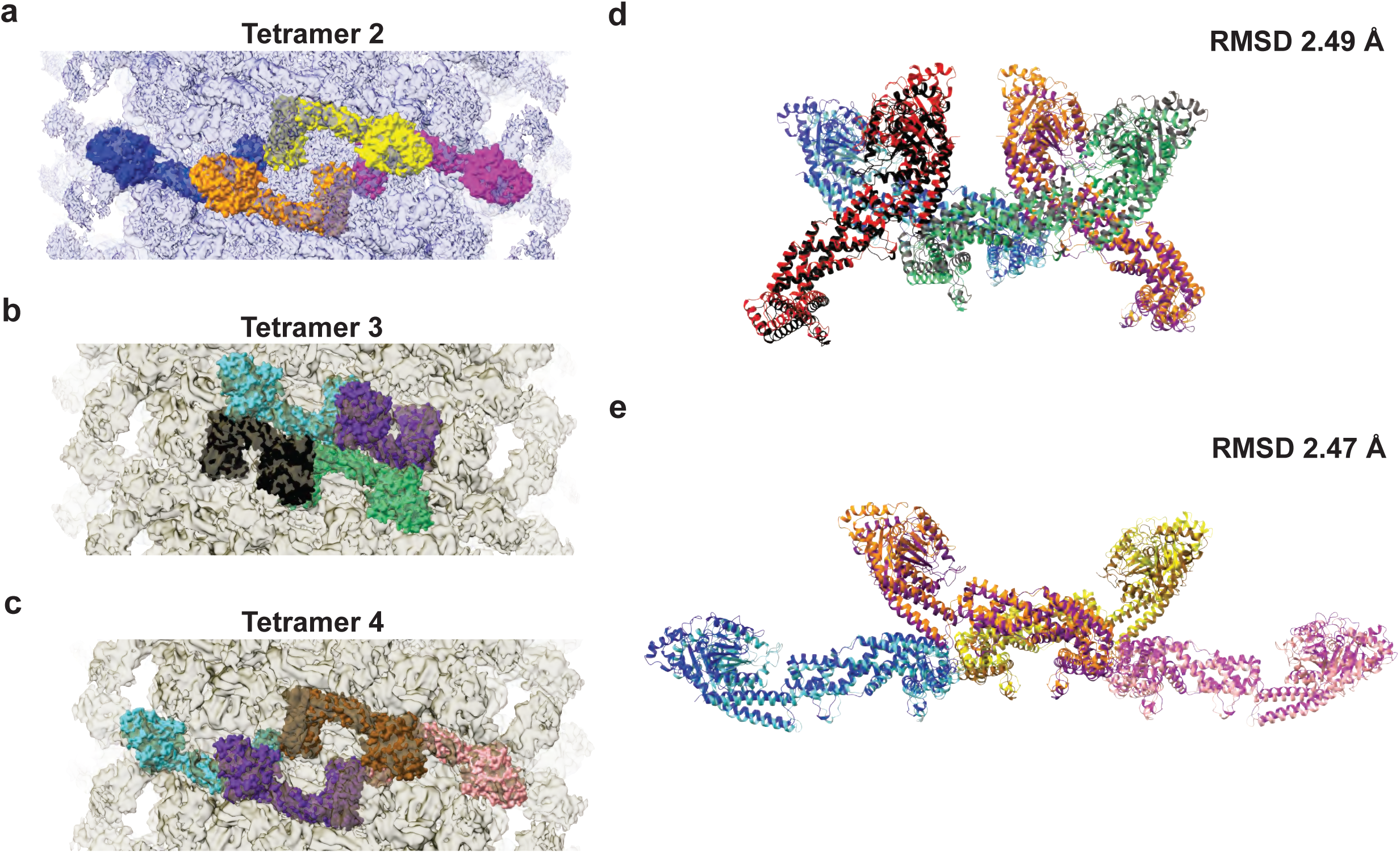
Description of four OPA1 tetramers used in MD simulations. **(a-c)** Three tetrameric subassemblies of the S-OPA1 polymer (tetramers 2, 3, and 4) were fitted into the corresponding density map, which is transparently visible. Each monomer is shown in surface representation and colored differently for clarity. Tetramer 2 **(a)** is extracted from the polymeric model in membrane-proximal conformation, while tetramers 3 **(b)** and 4 **(c)** are extracted from the polymeric model that represents the membrane-distal conformation of the S-OPA1 polymer. **(d, e)** Superimposition of the S-OPA1 tetramers assembled using different oligomerization interfaces. **(d)** Tetramers representing the conserved crisscross association of dynamin superfamily proteins. **(e)** The newly identified interface 7 mediates the formation of other tetrameric assemblies in the membrane-bound state. The root-mean-square deviation (RMSD) is calculated using the CLICK server.

**Supplementary Figure 2.**
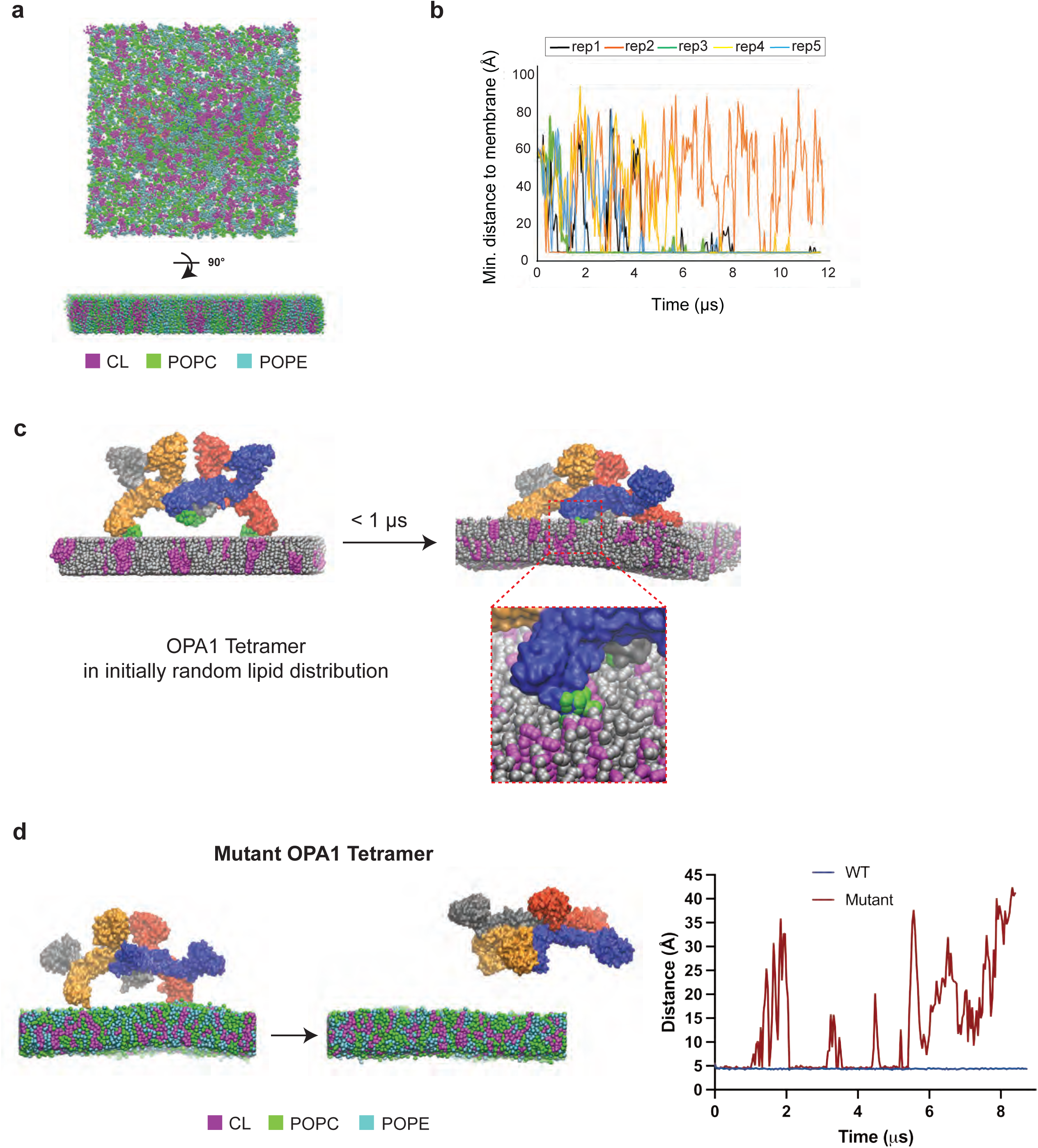
An overview of the MD simulation setup. **(a)** A representative image of the membrane patch used in CG MD simulations. The lipid molecules are shown in magenta (CL), green (POPC), and cyan (POPE). **(b)** The trajectories of OPA1-membrane interactions using the S-OPA1 starting model positioned ∼60 Å away from the membrane. The graph shows the minimal distance between the protein and membrane, calculated from five independent replicas of the simulation. **(c)** S-OPA1 tetramer binds to the membrane patch via the membrane-inserting loop (MIL) and docking regions in CG MD simulations. The subunits of the S-OPA1 tetramer are colored blue, yellow, orange, and gray. The MIL region is highlighted in green. S-OPA1 tetramers are positioned close to the membrane patch in the simulations. After a <1 μs simulation time, the tetramers rapidly formed charge-charge and hydrophobic interactions with the bilayer lipids and deformed the membrane patch. **(d)** The S-OPA1 tetramers containing mutations within the MIL and docking regions do not bind the membrane patch and remain in solution within the timescale of the simulations.

**Supplementary Figure 3.**
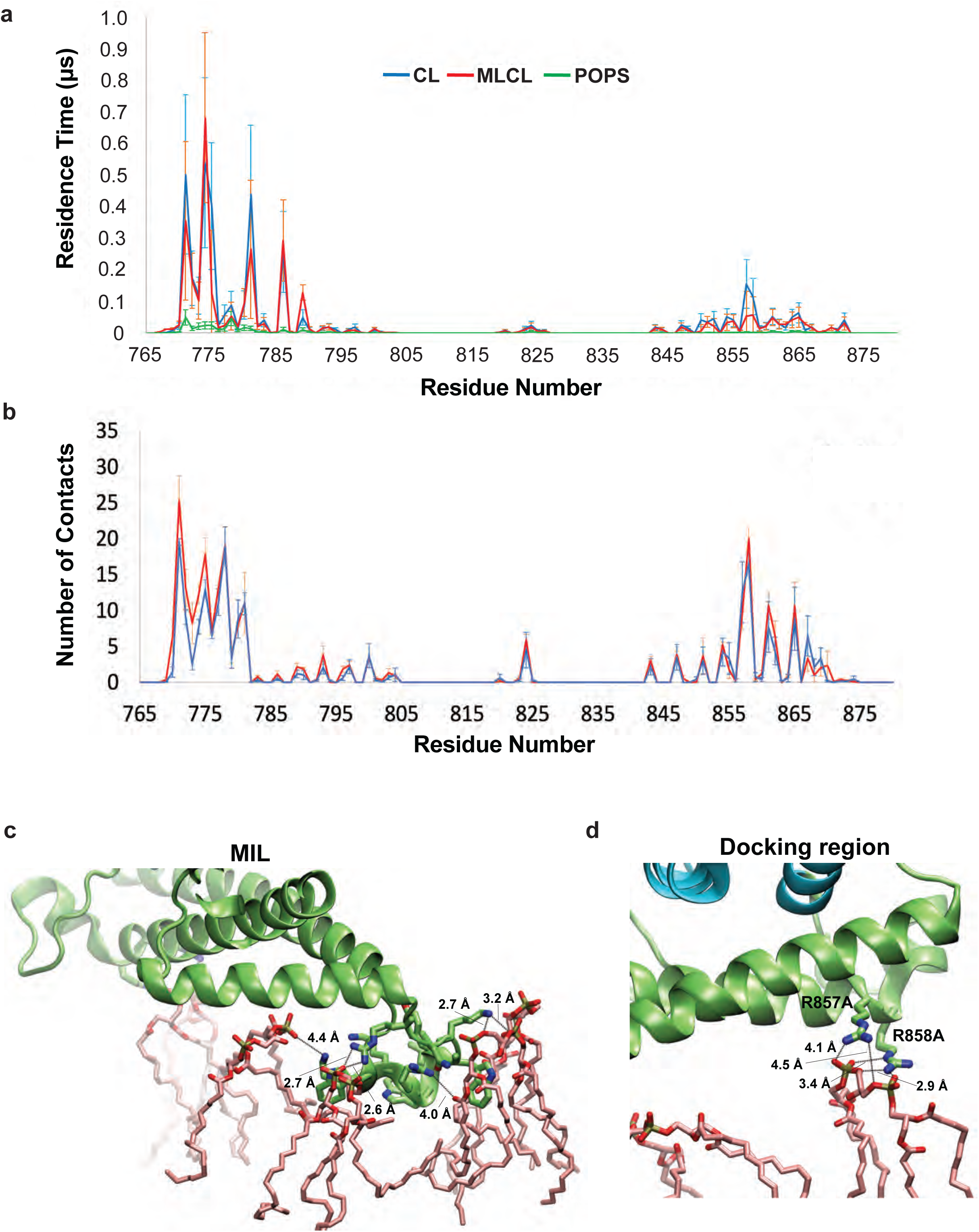
Comparison of residence times for CL and MLCL lipids in AA and CG MD simulations. **(a)** Residence times for contacts between S-OPA1 residues and CL (blue line), MLCL (red line), and POPS (green line) lipids in CG MD simulations. The data was averaged over four subunits in each tetramer and three replicas. **(b)** Average number of protein-lipid contacts calculated from three replicas of AA MD simulations using S-OPA1 tetramer and CL- and MLCL-enriched membranes. **(c, d)** Molecular interactions between S-OPA1 and CL headgroups. Close-up views of the two CL-binding motifs highlight key protein-lipid interactions in AA MD simulations. The paddle domain of S-OPA1 is shown in green, and CL molecules are colored in light pink. The distances between S-OPA1 residues and CL molecules are shown in Angstroms.

**Supplementary Figure 4.**
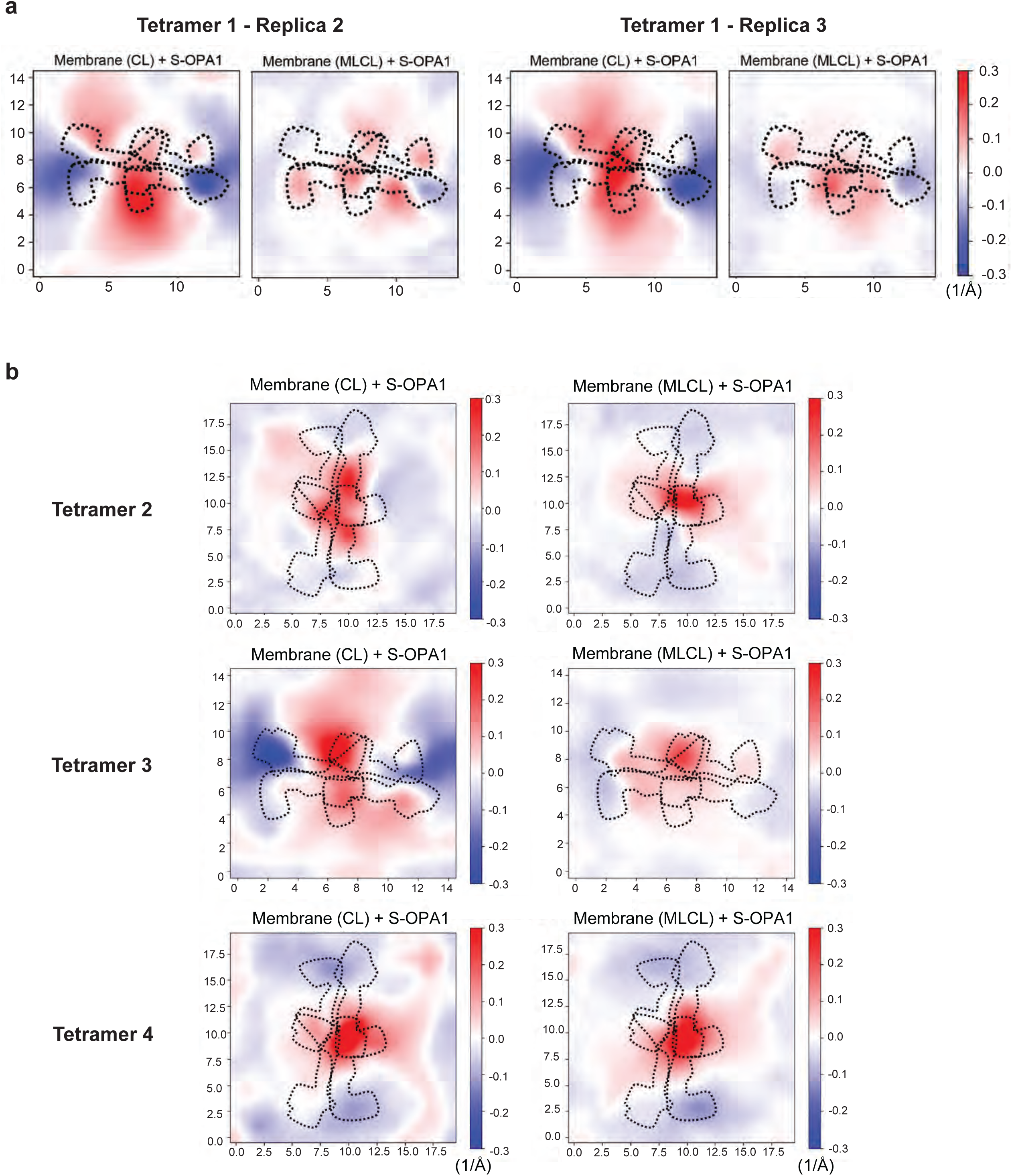
Membrane deformation analysis of S-OPA1 tetramer in CG MD simulations. Red and blue colors indicate membrane pulling and pushing in the direction of z, respectively. The x and y axes indicate the number of membrane tiles; each tile represents 15 Å. **(a)** Membrane deformation calculations are shown for two other independent replicas using S-OPA1 tetramer 1 and model membranes containing either CL (left) or MLCL (right). The membrane deformation activity of tetramer 1, particularly its ability to push down on the sides, is reduced in the presence of MLCL. **(b)** Average membrane deformation was calculated for S-OPA1 tetramers 2, 3, and 4. A comparison of CL- and MLCL-containing membranes indicates reduced membrane deformation in the presence of MLCL for tetramer 3. While the CG MD simulations with tetramers 1 and 3 display significant membrane bending with CL-containing membranes, tetramers 2 and 4 show no visible difference between the two membranes.

**Supplementary Figure 5.**
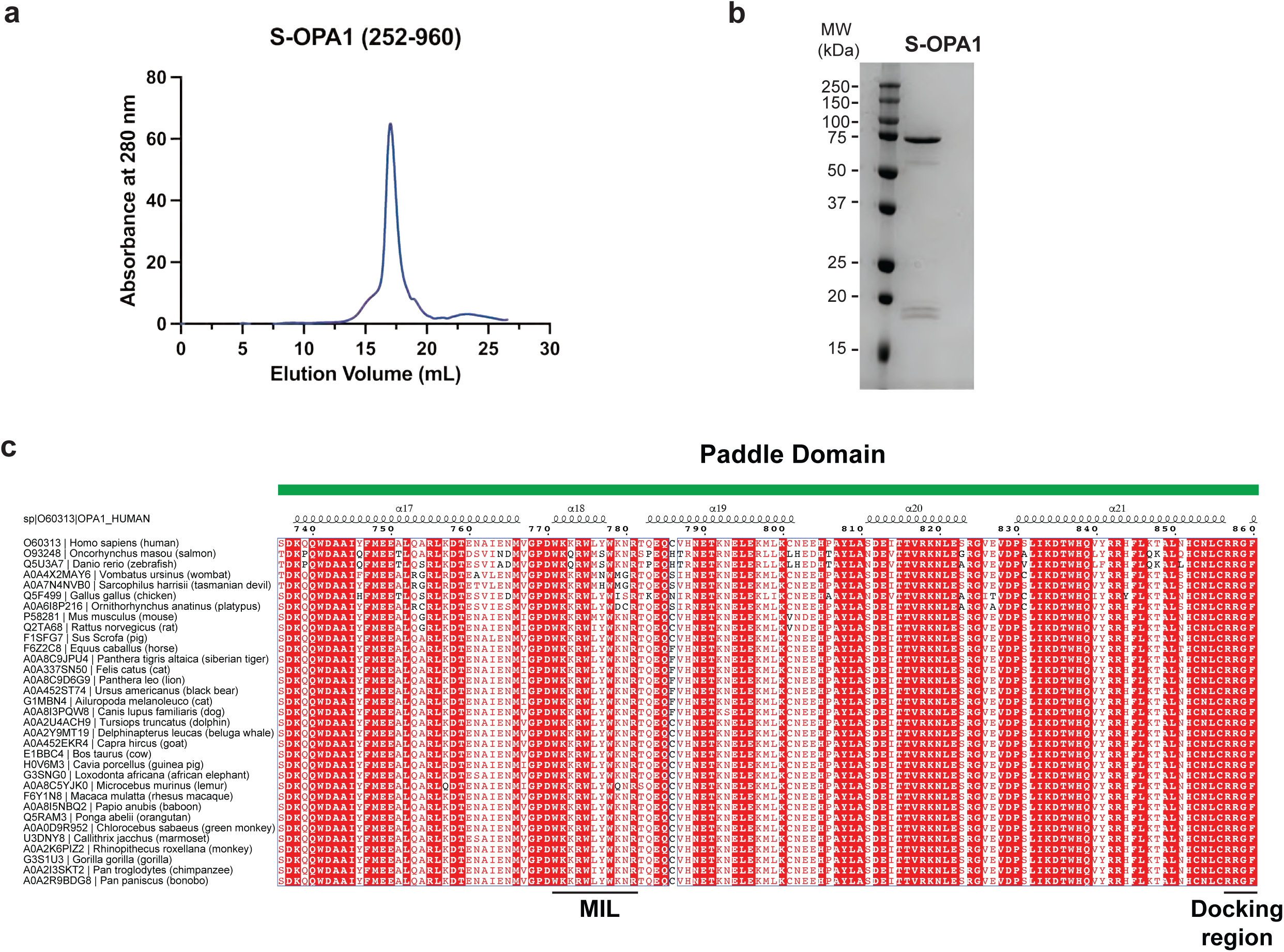
Biochemical characterization of human S-OPA1 construct and sequence alignment of the paddle domain. **(a)** A representative size-exclusion chromatography (SEC) profile of S-OPA1 WT and **(b)** SDS-PAGE of S-OPA1 protein following SEC. **(c)** The sequence alignment of paddle domain residues (736 to 860) demonstrates high sequence conservation across 33 species. Solid black lines below the sequence alignment indicate the boundaries of the membrane-inserting loop (MIL) and docking regions.

**Supplementary Figure 6.**
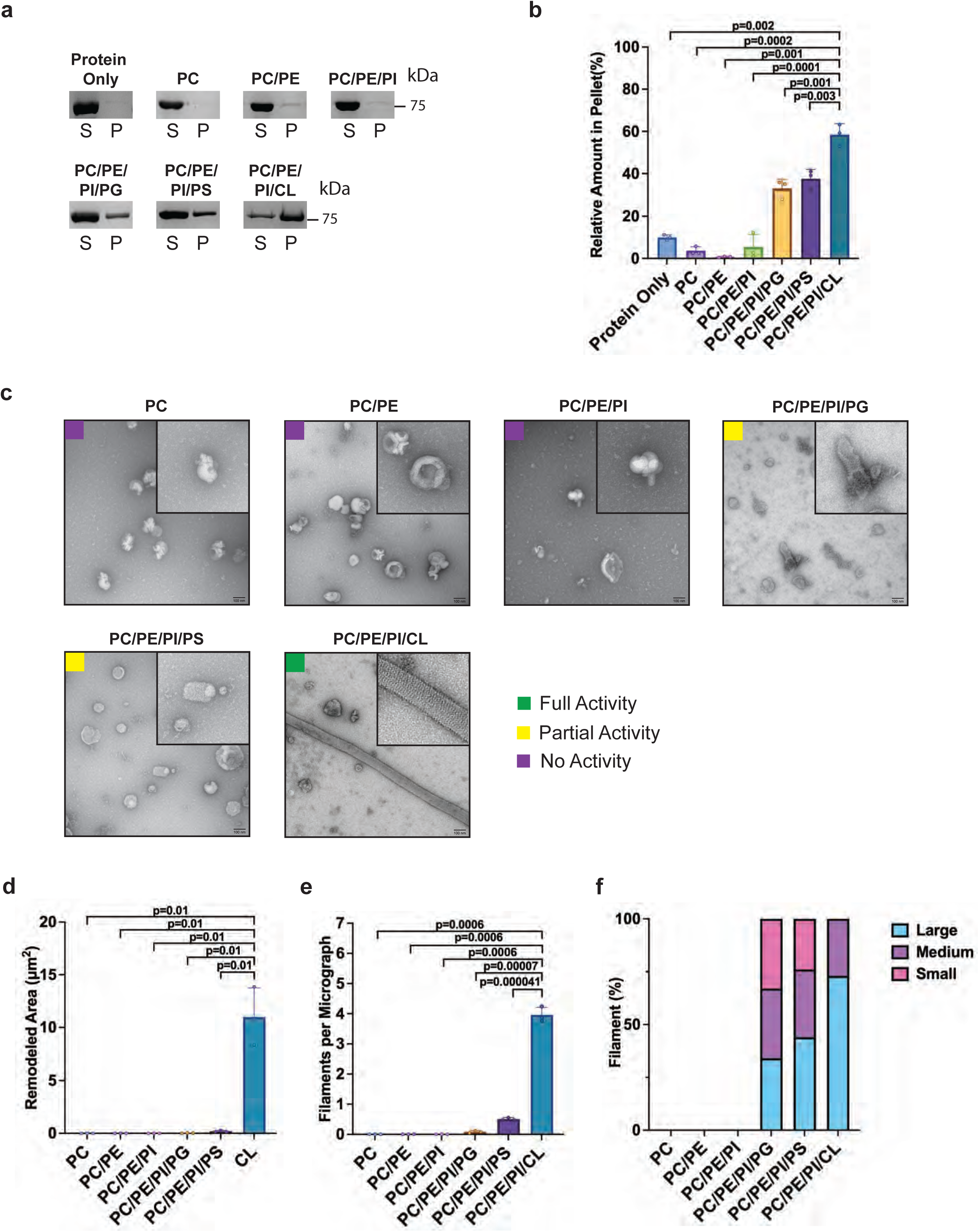
Membrane binding and remodeling experiments of S-OPA1 with various lipids. **(a)** Co-sedimentation assays of S-OPA1 WT using six different liposomes containing POPC, POPE, POPS, POPG, L-PI, and CL at various concentrations. The PC/PE/PI/CL liposomes contain 45% POPC, 22% POPE, 8% L-PI, and 25% CL; The PC/PE/PI/PS liposomes contain 45% POPC, 22% POPE, 8% L-PI, and 25% POPS; The PC/PE/PI/PG liposomes contain 45% POPC, 22% POPE, 8% L-PI, and 25% POPG; the PC/PE/PI liposomes contain 70% POPC, 22% POPE, and 8% L-PI; the PC/PE liposomes contains 78% POPC and 22% POPE; and the PC liposomes contain 100% POPC. Supernatant and pellet samples from the co-sedimentation assays were harvested after centrifugation and analyzed by SDS-PAGE. **(b)** The assays were performed in triplicate and gel images were quantified by ImageJ. The binding activity of S-OPA1 WT was quantified from pellet fractions, and statistical significance was assessed using a two-tailed unpaired Welch’s t-test based on *n*=3 independent experiments, comparing CL-containing liposomes to other liposomes with various compositions. *P*-values are represented on the graph, *and p*<0.05 is considered to be statistically significant. **(c)** Reconstitution assays were performed with the same liposomes as in **(a)**. The samples were incubated for ∼4 hours at room temperature and visualized by using negative-stain TEM. Scale bar is 100 nm. **(d)** Remodeled area (μm^2^) was calculated by measuring the surface area of membrane tubules. Each lipid composition was tested in three independent reconstitution assays, and negative stain TEM imaging was performed on each replicate. The total remodeled area was calculated per micrograph, and a total of 75 negative stain micrographs were analyzed per lipid composition. **(e)** S-OPA1 filaments were counted in each micrograph and plotted against the corresponding lipid composition. **(d, e)** Error bars represent the standard error of the mean (s.e.m.). Statistical significance was assessed using a two-tailed unpaired Welch’s *t*-test with *n*=3 independent experiments. *P*-values are annotated on the graph, and *p*<0.05 is considered statistically significant. **(f)** Quantification of filament morphology per liposome composition. Each remodeled filament was counted and assigned to one of the three classes: large, medium, or small. Filament sizes were defined as small at 0-0.01 μm^2^ (pink), medium for 0.01-0.05 μm^2^ (purple), and large if >0.05 μm^2^ (cyan). The percentage of each class is calculated per sample and is shown as a stacked bar graph.

**Supplementary Figure 7.**
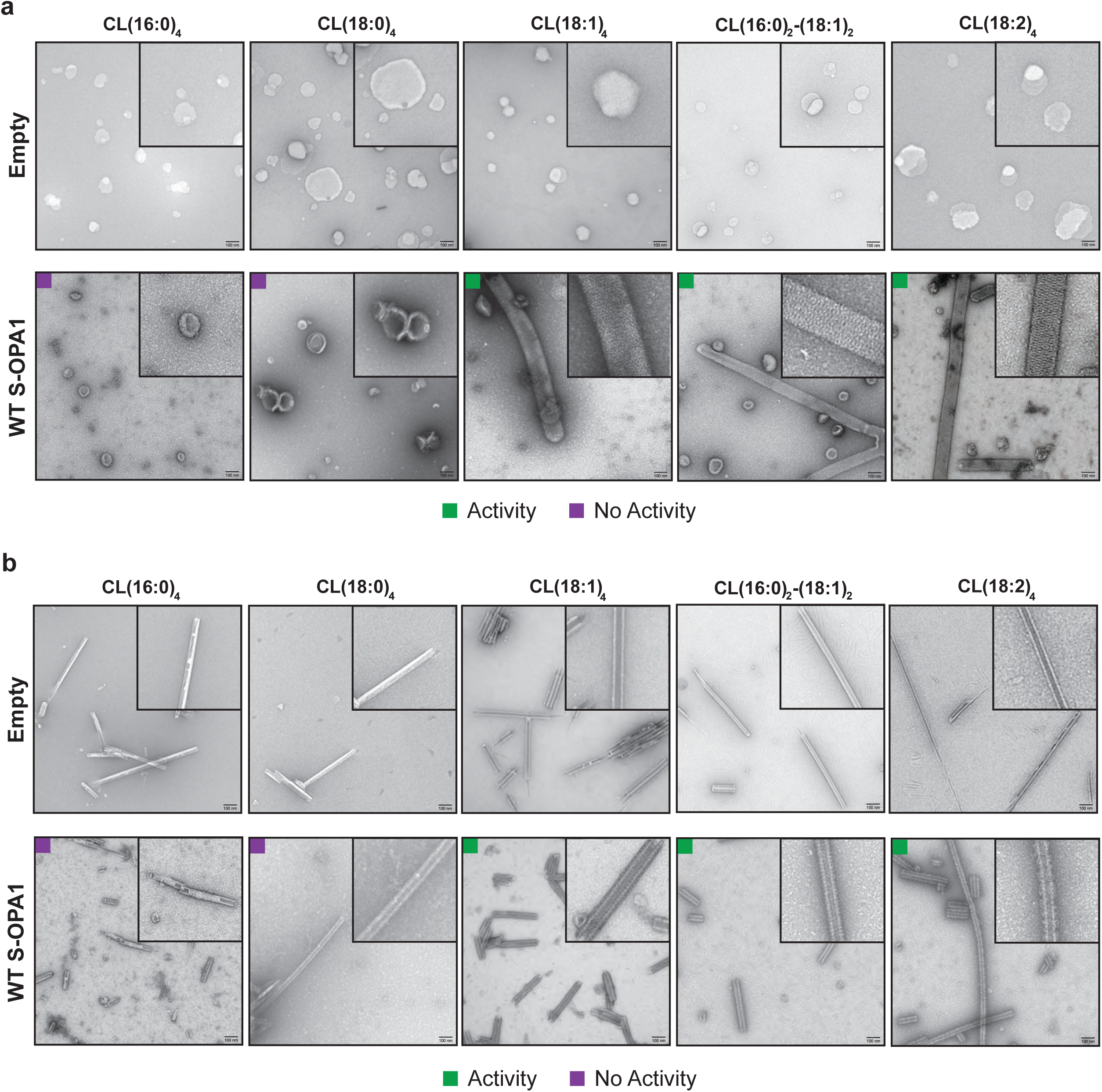
Negative-stain TEM images of liposomes and lipid nanotubes for different CL species. **(a)** Representative negative-stain TEM images show empty liposomes and membrane tubulation in the presence of S-OPA1 WT. All liposomes share a common composition of 45% POPC, 22% POPE, and 8% L-PI, with the remaining 25% composed of five different CL species: CL(16:0)_4_, CL(18:0)_4_, CL(18:1)_4_, CL(16:0)_2_-(18:1)_2_, and CL(18:2)_4._ Protein samples were able to bind and remodel membranes containing CL(18:1)_4_, CL(16:0)_2_-(18:1)_2_, and CL(18:2)_4_, whereas liposomes prepared with CL(16:0)_4_, and CL(18:0)_4_ impaired S-OPA1’s ability to bind and remodel lipid membranes. **(b)** Negative-stain TEM images of lipid nanotubes composed of five different CL species, CL(16:0)_4_, CL(18:0)_4_, CL(18:1)_4_, CL(16:0)_2_-(18:1)_2_, and CL(18:2)_4_, in the presence and absence of S-OPA1 WT. The nanotubes were composed of 90% Galactosyl(β) Ceramide and 10% of the respective CL species. Similar to above experiments, lipid nanotubes composed of CL(16:0)_4_, and CL(18:0)_4_ hindered S-OPA1-membrane interactions, while CL(18:1)_4_, CL(16:0)_2_-(18:1)_2_, and CL(18:2)_4_ nanotubes promoted the formation of S-OPA1 filaments on pre-curved membranes *in vitro*. Scale bars are 100 nm.

**Supplementary Figure 8.**
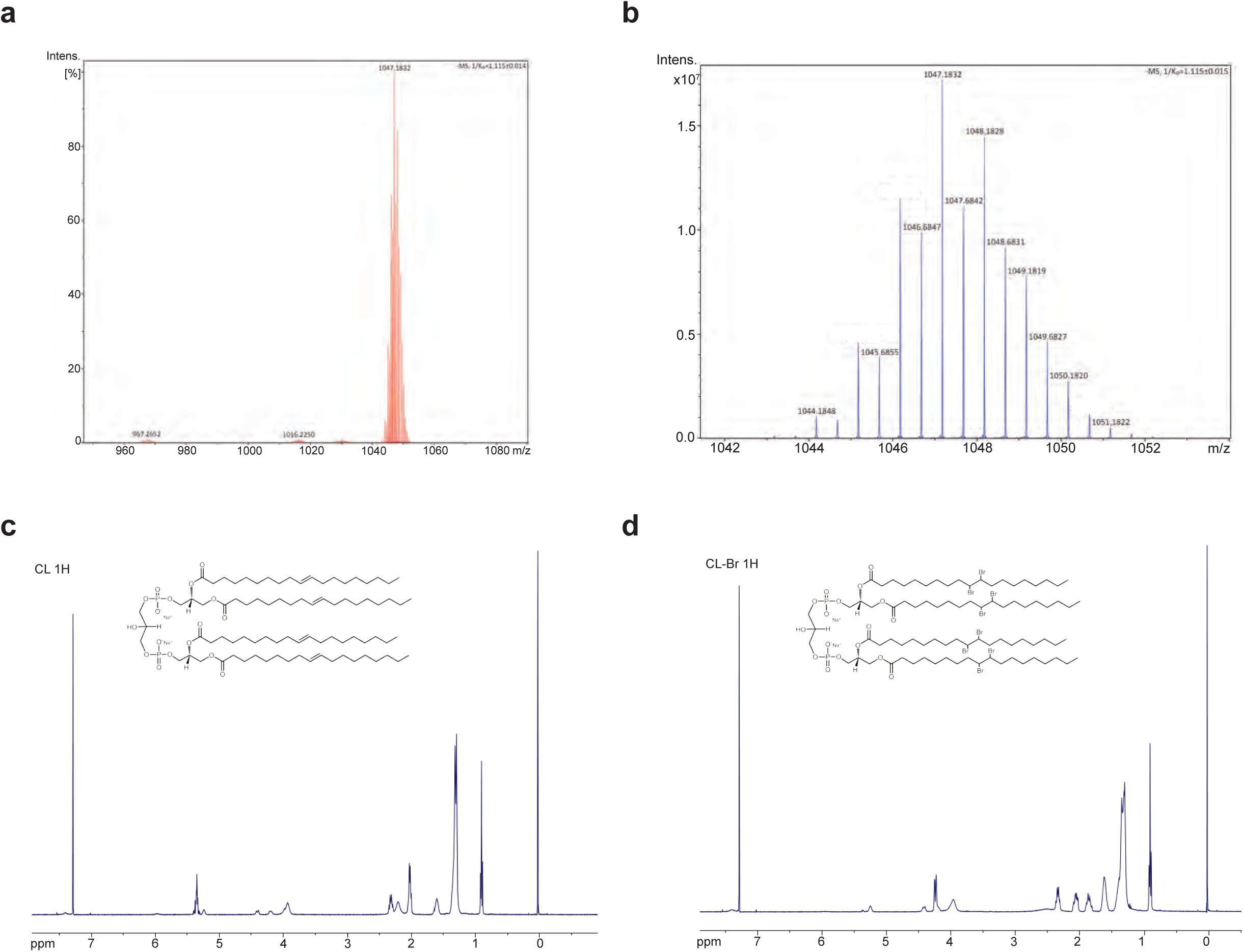
Validation of brominated cardiolipin. **(a)** Mass spectrum of brominated cardiolipin from 780 to 1280 mass to charge ratio (m/z). **(b)** Zoomed-in view of the mass spectrum from 1042 to 1052 m/z. **(c, d)** The brominated cardiolipin chemistry was validated by small ligand NMR. The NMR Spectrum of cardiolipin H^1^ **(c)** and brominated cardiolipin H^1^ **(d)**.

**Supplementary Figure 9.**
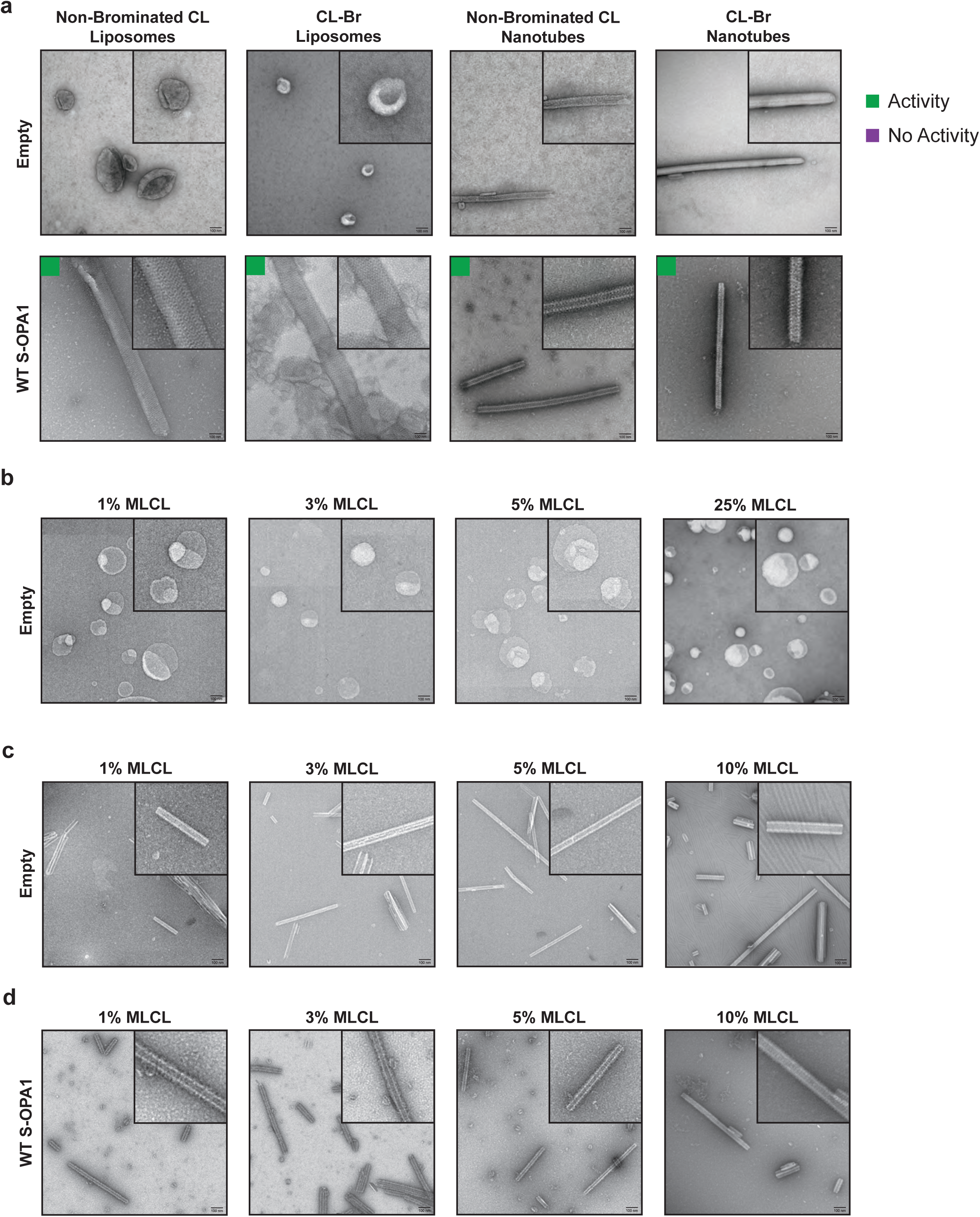
Negative-stain TEM images of various liposomes and lipid nanotubes. **(a)** Representative negative-stain TEM images of reconstitution assays show cylindrical and spherical liposomes in the presence and absence of S-OPA1 WT. All liposomes share a common composition of 45% POPC, 22% POPE, and 8% L-PI, with the remaining 25% composed of either native or brominated CL. Protein samples bind and form higher-order assemblies on brominated and native liposomes. Different molar concentrations of CL and MLCL were used to prepare various liposomes **(b)** and lipid nanotubes **(c)**. **(b)** Liposomes were prepared with varying CL:MLCL ratios, maintaining the same base lipid composition as in **(a)**. The CL:MLCL ratios used for liposome preparations are as follows: 24% CL and 1% MLCL; 22% CL and 3% MLCL; 20% CL and 5% MLCL; and 25% MLCL. **(c)** Similarly, the lipid nanotubes were composed of 90% Galactosyl(ß) Ceramide, and the remaining 10% consisted of the following CL:MLCL ratios: 9% CL and 1% MLCL; 7% CL and 3% MLCL; 5% CL and 5% MLCL; and 10% MLCL. **(b, c)** Negative-stain TEM micrographs show that increasing concentrations of MLCL do not change the morphology of liposomes and lipid nanotubes compared to the CL-enriched lipid bilayers. **(d)** Lipid nanotubes prepared with low molar concentrations of MLCL do not disrupt the formation of S-OPA1 filaments on pre-curved membranes *in vitro*. Scale bars are 100 nm.

**Supplementary Figure 10.**
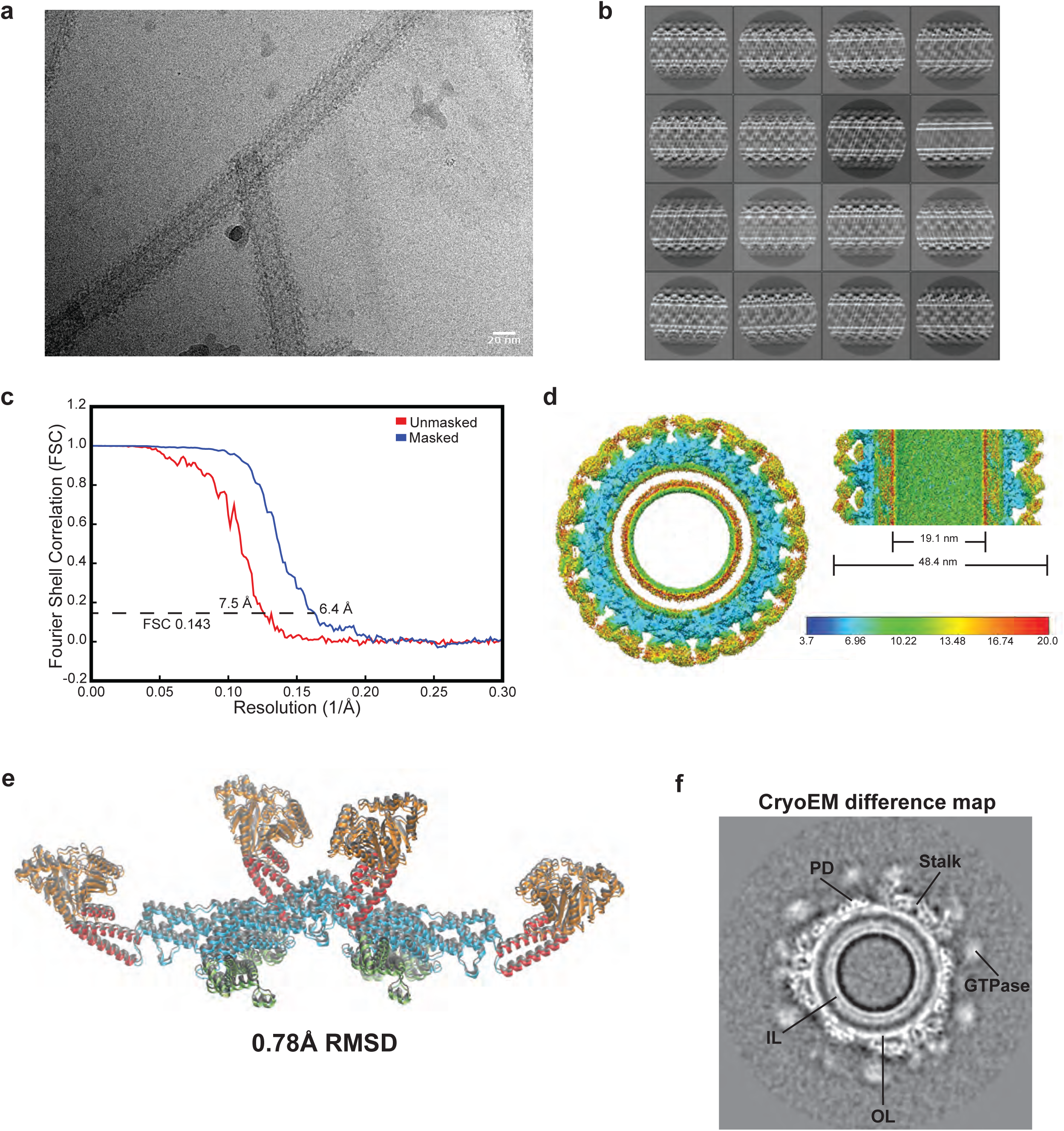
CryoEM imaging and image analysis of S-OPA1 assemblies bound to brominated liposomes. **(a)** Electron cryo-micrograph showing S-OPA1 filaments assembled on liposomes containing CL-Br. **(b)** Representative 2D class averages of S-OPA1 filament segments. **(c)** Gold-standard Fourier Shell Correlation (FSC) curve of the final density map. **(d)** Local resolution estimates for the cryoEM 3D reconstruction. Both horizontal and vertical slices through cryo-EM densities are shown. **(e)** S-OPA1 tetramer bound to brominated nanotubes (colored) superimposed with the tetrameric model bound to native nanotubes (gray) shows minimal structural differences between the two models. **(f)** A gray-scale slice of the difference map of S-OPA1 polymer bound to native and brominated liposomes shows CL enrichment in the outer leaflet. IL, Inner Leaflet; OL, Outer Leaflet; PD, Paddle Domain.

**Supplementary Figure 11.**
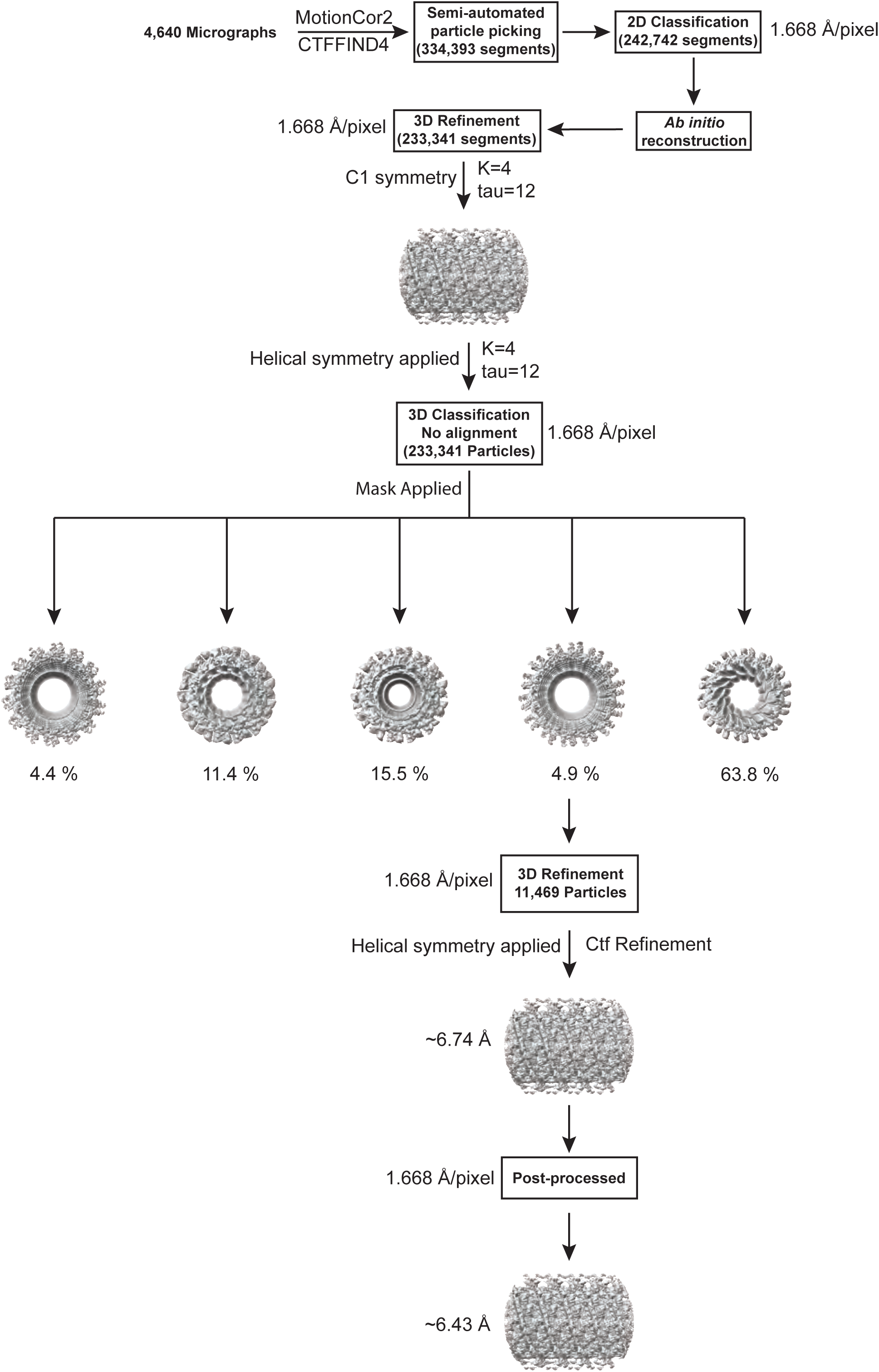
CryoEM data processing flowchart of S-OPA1 bound to brominated cardiolipin-containing membranes. Details of cryoEM data collection and image analysis are described in the methods section.

**Supplementary Table 1.** Cryo-EM data collection, refinement, and validation statistics for the two tetrameric S-OPA1 models.

|  | Human S-OPA1 bound to CL-Br membranes<br>(EMDB-43349) |  |
| --- | --- | --- |
| **Data collection and processing** |  |  |
| Microscope | FEI Titan Krios |  |
| Camera | Gatan K3 Summit |  |
| Magnification | 105,000x |  |
| Voltage (kV) | 300 |  |
| Electron exposure (e<sup>-</sup>/Å<sup>2</sup>) | 65 |  |
| Defocus (um) | 0.5 to 1.5 |  |
| Pixel Size (Å) | 0.834 |  |
| Symmetry imposed | Helical |  |
| Micrographs (no.) | 4,640 |  |
| Initial particle images (no.) | 233,341 |  |
| Final particle images (no.) | 11,469 |  |
| Map resolution (Å) | 6.4 |  |
| FSC threshold | 0.143 |  |
| Map resolution range (Å) | 4.8 to 8.1 | 4.8 to 8.2 |
| Models Generated (PDB code) | 8VLZ | 8VM4 |
| **Refinement** |  |  |
| Initial model used (PDB code) | 8CT1 |  |
| Model resolution (Å) | 6.4 |  |
| FSC threshold | 0.143 | 0.143 |
| Map sharpening B factor (Å<sup>2</sup>) | -316.754 |  |
| Model composition |  |  |
| Nonhydrogen atoms | 22,723 |  |
| Protein residues | 2,792 |  |
| B factors (Å<sup>2</sup>) – min |  |  |
| Protein | 112.87 | 131.06 |
| R.m.s deviations |  |  |
| Bond lengths (Å) | 0.002 | 0.002 |
| Bond angles (°) | 0.385 | 0.333 |
| **Validation** |  |  |
| MolProbity Score | 1.16 | 1.19 |
| Clashscore | 3.68 | 4.06 |
| Poor rotamers (%) | 0.0 | 0.0 |
| Ramachandran plot |  |  |
| Favored (%) | 99.57 | 99.57 |
| Allowed (%) | 0.43 | 0.43 |
| Disallowed (%) | 0.0 | 0.0 |

**Supplementary Table 2.**
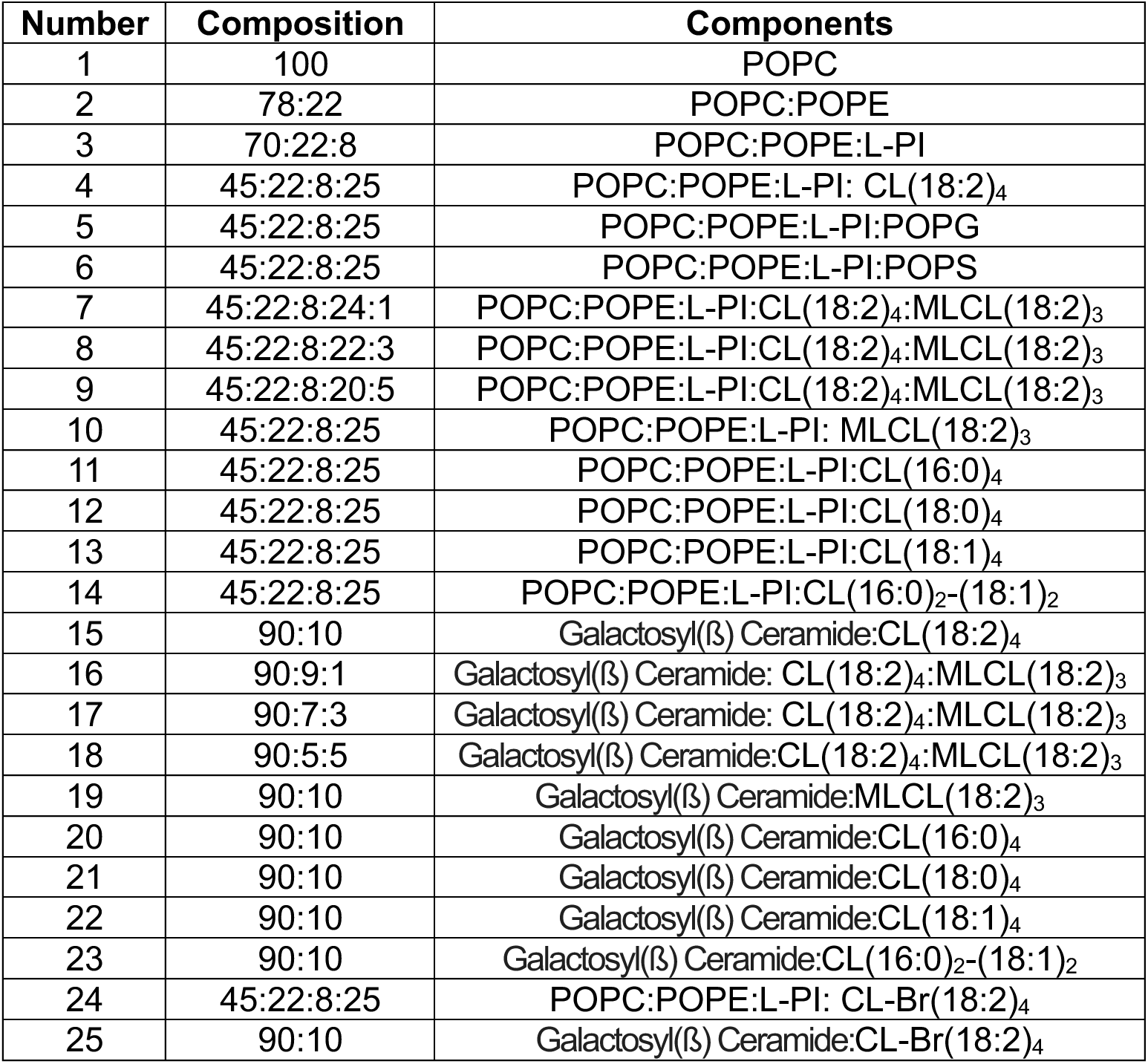
List of liposome compositions used in reconstitution assays, co-sedimentation experiments, and cryoEM imaging.

| Number | Composition | Components |
| --- | --- | --- |
| 1 | 100 | POPC |
| 2 | 78:22 | POPC:POPE |
| 3 | 70:22:8 | POPC:POPE:L-PI |
| 4 | 45:22:8:25 | POPC:POPE:L-PI: CL(18:2)<sub>4</sub> |
| 5 | 45:22:8:25 | POPC:POPE:L-PI:POPG |
| 6 | 45:22:8:25 | POPC:POPE:L-PI:POPS |
| 7 | 45:22:8:24:1 | POPC:POPE:L-PI:CL(18:2)<sub>4</sub>:MLCL(18:2)<sub>3</sub> |
| 8 | 45:22:8:22:3 | POPC:POPE:L-PI:CL(18:2)<sub>4</sub>:MLCL(18:2)<sub>3</sub> |
| 9 | 45:22:8:20:5 | POPC:POPE:L-PI:CL(18:2)<sub>4</sub>:MLCL(18:2)<sub>3</sub> |
| 10 | 45:22:8:25 | POPC:POPE:L-PI: MLCL(18:2)<sub>3</sub> |
| 11 | 45:22:8:25 | POPC:POPE:L-PI:CL(16:0)<sub>4</sub> |
| 12 | 45:22:8:25 | POPC:POPE:L-PI:CL(18:0)<sub>4</sub> |
| 13 | 45:22:8:25 | POPC:POPE:L-PI:CL(18:1)<sub>4</sub> |
| 14 | 45:22:8:25 | POPC:POPE:L-PI:CL(16:0)<sub>2</sub>-(18:1)<sub>2</sub> |
| 15 | 90:10 | Galactosyl(β) Ceramide:CL(18:2)<sub>4</sub> |
| 16 | 90:9:1 | Galactosyl(β) Ceramide: CL(18:2)<sub>4</sub>:MLCL(18:2)<sub>3</sub> |
| 17 | 90:7:3 | Galactosyl(β) Ceramide: CL(18:2)<sub>4</sub>:MLCL(18:2)<sub>3</sub> |
| 18 | 90:5:5 | Galactosyl(β) Ceramide:CL(18:2)<sub>4</sub>:MLCL(18:2)<sub>3</sub> |
| 19 | 90:10 | Galactosyl(β) Ceramide:MLCL(18:2)<sub>3</sub> |
| 20 | 90:10 | Galactosyl(β) Ceramide:CL(16:0)<sub>4</sub> |
| 21 | 90:10 | Galactosyl(β) Ceramide:CL(18:0)<sub>4</sub> |
| 22 | 90:10 | Galactosyl(β) Ceramide:CL(18:1)<sub>4</sub> |
| 23 | 90:10 | Galactosyl(β) Ceramide:CL(16:0)<sub>2</sub>-(18:1)<sub>2</sub> |
| 24 | 45:22:8:25 | POPC:POPE:L-PI: CL-Br(18:2)<sub>4</sub> |
| 25 | 90:10 | Galactosyl(β) Ceramide:CL-Br(18:2)<sub>4</sub> |

